## Supplementary Information for "A stochastic analysis of the interplay between antibiotic dose, mode of action, and bacterial competition in the evolution of antibiotic resistance"

### Contents


**A Bacterial population dynamics** S3
A.1 Competitive Lotka-Volterra-like population dynamics . . . . . . S4
A.2 Default parameter set . . . . . . S5

**B Survival probability of the resistant subpopulation during treatment** S6
B.1 Reviewing the general approach . . . . . . S6
B.2 Death competition . . . . . . S6
B.2.1 Biostatic antibiotic . . . . . . S6
B.2.2 Biocidal antibiotic . . . . . . S7
B.3 Birth competition . . . . . . S8
B.3.1 Biostatic antibiotic . . . . . . S8
B.3.2 Biocidal antibiotic . . . . . . S11

**C Predicting the antibiotic concentration minimizing resistant strain survival** S12
C.1 Scenario: Biocidal drug, birth competition . . . . . . S12
C.2 Scenario: Biocidal drug, death competition . . . . . . S12
C.3 Scenario: Biostatic drug, death competition . . . . . . S13

**D Resistant population size at end of treatment** S15
D.1 Frequency of resistant strain at the end of treatment . . . . . . S17

**E Mean carriage time of the resistant population after treatment** S18
E.1 Reviewing the method of timescale separation in Lotka-Volterra systems . . . . . . S18
E.2 Death competition . . . . . . S18
E.3 Birth competition . . . . . . S20
E.4 Comparison between deterministic and stochastic prediction . . . . . . S20

**F De-novo emergence of antibiotic resistance** S22

**G Alternative parameterization of the model** S26
G.1 Survival probability . . . . . . S26
G.2 Size at the end of treatment . . . . . . S27
G.3 Carriage time of the resistant strain . . . . . . S27

**H Alternative interaction of antibiotic and density-dependent processes** S29
H.1 Population dynamics . . . . . . S29
H.2 Survival probability . . . . . . S30

**I Alternative antibiotic response curve parameterizations** S32
I.1 Survival probability . . . . . . S32
I.2 Size at the end of treatment . . . . . . S35

**J Alternative model with an explicit immune response** S37
J.1 Model description . . . . . . S37
J.2 Sample trajectories . . . . . . S37
J.3 Survival probability . . . . . . S39

**K Numerical simulations** S41

### A Bacterial population dynamics

We present the mathematical details of a competitive Lotka-Volterra-like model, as described in the main text. We study two models that differ in the way how bacterial density affects the population dynamics. Density either affects the birth rate of cells (birth competition), thus entering in the term birth rate $\lambda_j(x_S, x_R)$, or it affects the death rate (death competition), changing $\mu_j(x_S, x_R)$. In addition, we examine the effects of two modes of action of antibiotic drugs. Biostatic drugs reduce the replication rate of bacterial cells, therefore reducing $\lambda_j(x_S, x_R)$, while biocidal drugs increase the death rate of bacteria, thus adding to $\mu_j(x_S, x_R)$. For clarity, we restate Table 1 from the main text that lists the birth and death rates for the four different modeling scenarios (Table A.1).

For sufficiently large population sizes, the dynamics of the population are well described by ordinary differential equations. For example, in our main model with birth competition and a biocidal antibiotic drug, the dynamics of the sensitive population density, denoted by $x_S(t)$, in the absence of the resistant strain is given by

$$\frac{dx_S(t)}{dt} = (\lambda_S(x_S(t), 0) - \mu_S(x_S(t), 0))x_S(t) = \left(\max\left(\beta_S - \gamma_S x_S(t), 0\right) - \delta_S - \alpha_S(c)\right) x_S(t) \,. \tag{A.1}$$

For population densities below the carrying capacity, so that the birth rate $\lambda(x_S, 0)$ is positive, the ordinary differential equation has the following solution, which can be found by any smybolic programming language, e.g. *Mathematica* (files can be found at `https://github.com/pczuppon/AMRWithinHost`)

$$x_S(t) = \frac{x_S(0)}{\frac{\gamma_S x_S(0)}{(\beta_S - \delta_S - \alpha_S(c))} + \left(1 - \frac{\gamma_S x_S(0)}{(\beta_S - \delta_S - \alpha_S(c))}\right) e^{-(\beta_S - \delta_S - \alpha_S(c))t}} \,. \tag{A.2}$$

The case of a biocidal drug in a model where density affects the death rate is treated analogously. In the remaining scenarios for bacteriostatic drugs, the solution cannot be stated in this general form. Rather, it needs to be split into different cases, depending on whether the antibiotic concentration is large enough so that the antibiotic effect, $\alpha_S(c)$, is larger than the birth rate $\beta_S - \gamma_S x_S(t)$ or not. A comparison of the different scenarios and their effect on the deterministic population dynamics of the sensitive population during treatment is shown in Fig. A.1. For antibiotic concentrations far below the minimal inhibitory concentration (MIC) of the sensitive strain, the scenarios do not differ. If the antibiotic concentration is slightly below the MIC, the only scenario that differs from the others is a biostatic treatment with birth competition (Fig. A.1 middle panel). The population size approaches the new, lower stationary population size slower than the other three scenarios. For

| | | **birth competition** | **death competition** |
|---|---|---|---|
| **biostatic** | $\lambda_j(x_S, x_R)$ | $\max\left(\beta_j - \gamma_j(x_S + x_R) - \alpha_j(c), 0\right)$ | $\max\left(\beta_j - \alpha_j(c), 0\right)$ |
| | $\mu_j(x_S, x_R)$ | $\delta_j$ | $\delta_j + \gamma_j(x_S + x_R)$ |
| **biocidal** | $\lambda_j(x_S, x_R)$ | $\max\left(\beta_j - \gamma_j(x_S + x_R), 0\right)$ | $\beta_j$ |
| | $\mu_j(x_S, x_R)$ | $\delta_j + \alpha_j(c)$ | $\delta_j + \gamma_j(x_S + x_R) + \alpha_j(c)$ |

Table A.1: **Birth and death rates in the four studied scenarios.** The overall birth and death rates, denoted by $\lambda_j$ and $\mu_j$, are composed by the birth, death and competition processes that occur at rates $\beta_j$, $\delta_j$ and $\gamma_j$, respectively. Additionally, the effect of antibiotics, denoted by $\alpha_j$, is affecting either the birth or the death rate, depending on the type of drug administered. The index $j$ indicates the strain-specificity, $j = S$ or $j = R$ for antibiotic-sensitive or -resistant cells.

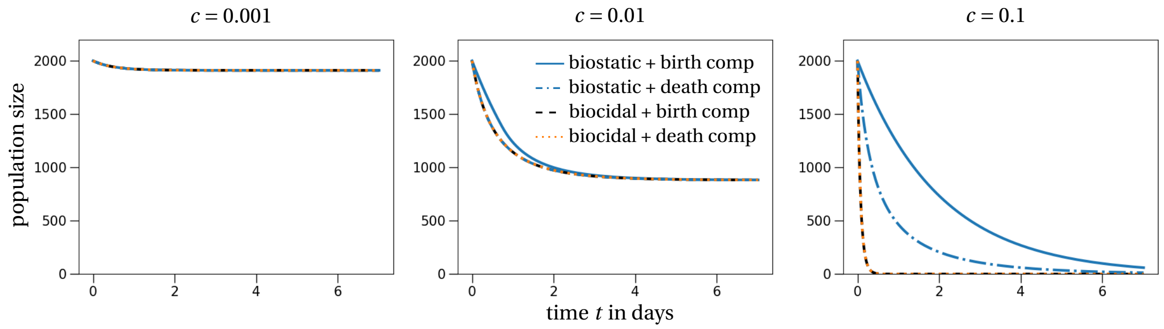


Figure A.1: **Deterministic dynamics of the sensitive strain under antibiotic treatment.** The sensitive population sizes decrease at the same speed for antibiotic concentrations far below the MIC of the sensitive strain (left panel). Increasing the antibiotic concentration, we find that the population size decreases fastest in the models under biocidal treatment, no matter the modeling details regarding the density-dependent effect. We observe the slowest decrease for biostatic treatment and birth competition. The parameters are set according to our default parameter set from the main text as stated in Table A.2.

concentrations larger than the MIC values, the scenarios differ more markedly. In the two scenarios with biocidal treatment the population sizes decrease at the same speed and faster than for biostatic drugs. For biostatic treatment with birth competition, the population size decreases the slowest. The exponential decline rates, or negative exponential growth rates, of the different scenarios for high antibiotic concentrations explain this observation (from largest to lowest): biocidal and death competition = $\beta_S - \delta_S - \alpha_S(c) - \gamma x_S(t)$; biocidal and birth competition = $\max(\beta_S - \gamma x_S(t), 0) - \delta_S - \alpha_S(c)$; biostatic and death competition = $\max(\beta_S - \alpha_S(c), 0) - \delta_S - \gamma x_S(t)$; biostatic and birth competition = $\max(\beta_S - \gamma x_S(t) - \alpha_S(c), 0) - \delta_S$. Note that because of the sharp drop in sensitive population size in the two biocidal scenarios, the population density does not substantially affect the decline rate overall, which makes the trajectories overlap.

This general pattern is explained by a saturation of the effect of biostatic drugs due to the interaction with the maximum in the growth rate $\lambda_S$. In contrast, the effect of biocidal drugs attains its limiting effect at much higher concentrations, which explains why the population sizes decrease faster under biocidal treatment than under biostatic treatment. The difference between the two biostatic treatment scenarios is more subtle. In this case the different decrease rates are explained by the effect of density regulation. For birth competition, depending on the strength of the antibiotic effect, density-dependent regulation may just play a negligible role as seen for large antibiotic concentrations. In this case the growth rate $\lambda_S$ is zero independently of the bacterial density because the antibiotic drug is suppressing any bacterial replication. Therefore, the model with birth competition exhibits a slower population decrease than the model with death competition.

#### A.1 Competitive Lotka-Volterra-like population dynamics

The general model dynamics, including both the sensitive and resistant strains, are given by a competitive Lotka-Volterra-like model. It is not exactly a Lotka-Volterra model because of the maxima in the birth rates. For example, in our default scenario with birth competition and biocidal treatment, the corresponding system of ordinary differential equations is the following:

$$\begin{aligned}\frac{dx_S(t)}{dt} &= x_S(t)\left(\max\left(\beta_S - \gamma_S\left(x_S(t)+x_R(t)\right),0\right) - \delta_S - \alpha_S(c)\right),\\ \frac{dx_R(t)}{dt} &= x_R(t)\left(\max\left(\beta_R - \gamma_R\left(x_S(t)+x_R(t)\right),0\right) - \delta_R - \alpha_R(c)\right).\end{aligned} \tag{A.3}$$

#### A.2 Default parameter set

If not stated otherwise, we use the following default parameter set throughout all of the figures.

| **Biological meaning** | **Parameter** | **Default value** |
|---|---|---|
| birth rate sensitive strain | $\beta_S$ | 2.5 |
| birth rate resistant strain | $\beta_R$ | 2.25 |
| death rate sensitive strain | $\delta_S$ | 0.5 |
| death rate resistant strain | $\delta_R$ | 0.5 |
| competition rate | $\gamma$ | 1.0 |
| order of the carrying capacity | $K$ | 1,000 |
| minimal inhibitory concentration sensitive strain | $\text{mic}_S$ | 0.017 |
| maximal growth rate of strain $j$ | $\psi_{j,\max}$ | $\beta_j - \delta_j$ |
| maximal decline rate of strain $j$ under antibiotic treatment | $\psi_{j,\min}$ | $-156 \times \log(10)$ |
| steepness of antibiotic response curve | $\kappa$ | 1.1 |
| antibiotic concentration | $c$ | varies |
| treatment duration | $\tau$ | 7 (days) |
| initial density of sensitive cells | $x_S(0)$ | $(\beta_S - \delta_S)/\gamma$ |
| initial density of resistant cells | $x_R(0)$ | 1/K |
| initial number of sensitive cells | $X_S(0)$ | $(\beta_S - \delta_S)K/\gamma$ |
| initial number of resistant cells | $X_R(0)$ | 1 |
| mutation probability | $\eta$ | 0 (varied in Section F) |

Table A.2: **Default parameter set.**

### B Survival probability of the resistant subpopulation during treatment

We compute the probability that the resistant strain survives treatment. We use continuous-time branching processes in a varying environment to calculate this probability. Here, the environment is given by the declining sensitive subpopulation that affects the competition terms of the population dynamics. We start by reviewing the mathematical details of this method and then proceed to study all the different scenarios.

#### B.1 Reviewing the general approach

The general solution to the problem of survival of a branching process in a time-varying environment has been obtained by Kendall (1948). Here, we follow the notation and exposition from Uecker and Hermisson (2011) who studied this problem under different biologically relevant situations. We briefly recall the main assumptions and results that are necessary for our subsequent analysis.

The continuous-time branching process describes a population of cells, in our case the resistant strain, with birth rate $\lambda_R(x_S, x_R)$ and death rate $\mu_R(x_S, x_R)$. The deterministic population dynamics of the sensitive strain define the (time-varying) environment and are described by the growth rate $r(t) = \lambda_S(x_S(t), x_R(t)) - \mu_S(x_S(t), x_R(t))$. Using these definitions the selective advantage of the resistant strain during treatment is defined by

$$s(t) = \lambda_R(x_S(t), x_R(t)) - \mu_R(x_S(t), x_R(t)) - r(t)\,. \tag{B.1}$$

These are the ingredients of Eq. (8) in Uecker and Hermisson (2011). The survival probability of the mutant type (resistant strain) until time $\tau$, denoted $\varphi(\tau)$, if initially there are $n_0$ mutant cells, then reads (Eq. (13) in Uecker and Hermisson (2011)):

$$\varphi(\tau) = 1 - \left(1 - \frac{1}{1 + \int_0^\tau \mu_R(x_S(t), x_R(t)) \exp\left(\int_0^t (\lambda_R(x_S(t'), x_R(t')) - \mu_R(x_S(t'), x_R(t')))dt'\right) dt}\right)^{n_0}, \tag{B.2}$$

which for $n_0 = 1$ simplifies to (Eq. (16) in Uecker and Hermisson (2011))

$$\varphi(\tau) = \frac{1}{1 + \int_0^\tau \mu_R(x_S(t), x_R(t)) \exp\left(\int_0^t (\lambda_R(x_S(t'), x_R(t')) - \mu_R(x_S(t'), x_R(t')))dt'\right) dt}\,. \tag{B.3}$$

We now analyze this establishment probability for biostatic and biocidal antibiotics and under different forms of density-regulation.

#### B.2 Death competition

In this section, we derive the survival probability in the model where bacterial density increases the death rates (Table A.1, right column).

##### *B.2.1 Biostatic antibiotic*

Biostatic antibiotics suppress the reproduction of bacteria and therefore reduce their birth rate in our model. Following the notation introduced in the previous section, we write the birth rate of a mutant as

$$\lambda_R(x_S(t), x_R(t)) = \max\left(\beta_R - a_R(c), 0\right) \tag{B.4}$$

and the death rate as

$$\mu_R(x_S(t), x_R(t)) = \delta_R + \gamma_R x_S(t). \tag{B.5}$$

Note that the resistant subpopulation does not affect the competition term. This is an underlying assumption of our branching process approximation. We assume that the resistant subpopulation is small enough that it does not affect the relevant dynamics during the establishment phase.

We denote the time-dependent growth rate of the sensitive strain by

$$r(t) = \max(\beta_S - \alpha_S(c), 0) - \delta_S - \gamma_S x_S(t) \tag{B.6}$$

and we define the selective advantage of resistant cells under treatment by

$$\begin{aligned} s(t) &= \lambda(x_S(t), x_R(t)) - \mu(x_S(t), x_R(t)) - r(t) \\ &= \max(\beta_R - \alpha_R(c), 0) - \max(\beta_S - \alpha_S(c), 0) - \left(\delta_R - \delta_S + (\gamma_R - \gamma_S) x_S(t)\right) . \end{aligned} \tag{B.7}$$

Then, the survival probability until time $\tau$ of the resistant strain under a biostatic antibiotic, $\varphi_{\text{bs}}(\tau)$, can be computed numerically by Eq. (B.3). Unfortunately, an analytical solution is not accessible in this general case. If we assume that the competitiveness of the two strains is equal, $\gamma_R = \gamma_S = \gamma$, analytical progress is possible because the selective advantage $s(t)$ becomes time-independent. Furthermore, if we denote $\rho_k = \max(\beta_k - \alpha_k, 0) - \delta_k$ for $k$ being the strain type (either $S$ or $R$) and suppress the dependency of $\alpha_k$ on the antibiotic concentration $c$, we find

$$\varphi_{\text{bs}}^{\text{d}}(\tau) = \begin{cases} \frac{s\rho_S\rho_R}{s\rho_R\rho_S + x_S(0)\gamma\rho_R(\delta_R+\rho_S)(1-e^{-s\tau}) - \delta_R s(\gamma x_S(0) - \rho_S)(1-e^{-\rho_R\tau})}, & \text{if } \beta_R > \alpha_R \\ \frac{\rho_S e^{-\delta_R\tau}}{\rho_S - \gamma x_S(0)(1-e^{\rho_S\tau})}, & \text{else.} \end{cases} \tag{B.8}$$

For $\tau$ tending to infinity we find the establishment probability of the resistant strain which simplifies to

$$\begin{aligned} \varphi_{\text{bs}}^{\text{d}}(\infty) &= \max\left(0, \frac{s\rho_S\rho_R}{s\rho_R\rho_S + x_S(0)\gamma\rho_R(\delta_R+\rho_S) - \delta_R s(\gamma x_S(0) - \rho_S)}\right) \\ &= \max\left(0, \frac{1}{1 + \frac{\delta_R}{\rho_R}\left(\frac{\gamma x_S(0)}{s} + 1\right) + \frac{\gamma x_S(0)}{s}}\right) . \end{aligned} \tag{B.9}$$

The calculation to arrive at these formulas are summarized in a *Mathematica* notebook that is available at `https://github.com/pczuppon/AMRWithinHost`.

##### *B.2.2 Biocidal antibiotic*

Biocidal antibiotics actively kill bacteria. Therefore, the death rate is increased under such treatment. In this scenario the birth rate of a resistant cell is

$$\lambda_R(x_S(t), x_R(t)) = \beta_R \tag{B.10}$$

and the death rate is

$$\mu_R(x_S(t), x_R(t)) = \delta_R + \alpha_R(c) + \gamma_R x_S(t). \tag{B.11}$$

Repeating the analysis from above, i.e., assuming equal competition parameters $\gamma_R = \gamma_S = \gamma$ and setting $\rho_k = \beta_k - \delta_k - \alpha_k$ and $s = \rho_R - \rho_S$, the survival probability until time $\tau$ of a resistant strain under a biocidal antibiotic, $\varphi_{\text{bc}}(\tau)$, is given as

$$\varphi_{\text{bc}}^{\text{d}}(\tau) = \max\left(0, \frac{s\rho_S\rho_R}{s\rho_S\rho_R + \gamma x_S(0)\rho_R(\delta_R + \alpha_R + \rho_S)(1-e^{-s\tau}) - s(\gamma x_S(0) - \rho_S)(\delta_R + \alpha_R)(1-e^{-\rho_R\tau})}\right) . \tag{B.12}$$

Letting $\tau$ tend to infinity, this simplifies to

$$\begin{aligned}\varphi^{\mathrm{d}}_{\mathrm{bc}}(\infty) &= \max\left(0, \frac{s\rho_S\rho_R}{s\rho_S\rho_R + \gamma x_S(0)\rho_R(\delta_R+\alpha_R+\rho_S) - s(\gamma x_S(0)-\rho_S)(\delta_R+\alpha_R)}\right)\\ &= \max\left(0, \frac{1}{1+\frac{\delta_R+\alpha_R}{\rho_R}\left(\frac{\gamma x_S(0)}{s}+1\right)+\frac{\gamma x_S(0)}{s}}\right).\end{aligned} \tag{B.13}$$

#### B.3 Birth competition

In this section we outline the derivation of survival probabilities in the model where the pathogen density reduces the birth rate (Table A.1, left column).

##### *B.3.1 Biostatic antibiotic*

We again start with the case of treatment with a biostatic drug. Following the notation introduced in the previous sections, we write the birth rate of a resistant cell as

$$\lambda_R(x_S(t), x_R(t)) = \max\left(\beta_R - \gamma_R x_S(t) - \alpha_R(c), 0\right) \tag{B.14}$$

and the death rate as

$$\mu_R(x_S(t), x_R(t)) = \delta_R. \tag{B.15}$$

The time-dependent growth rate of the sensitive strain is given by

$$r(t) = \max\left(\beta_S - \alpha_S(c) - \gamma_S x_S(t), 0\right) - \delta_S. \tag{B.16}$$

The growth rates, excluding the terms corresponding to density regulation, are denoted by $\rho_k = \max(\beta_k - \alpha_k, 0) - \delta_k$.

For the analysis of the survival probability of the resistant strain during antibiotic treatment, we need to split up the analysis according to different parameter regions defined by the antibiotic concentration (treatment duration denoted by $\tau$): (i) low antibiotic concentration ($\alpha_S(c) < \beta_S - \gamma x_S(t)$ for all $t \le \tau$); (ii) intermediate antibiotic concentration ($\alpha_S(c) > \beta_S - \gamma x_S(t)$ initially and then switches sign for some $T < \tau$); (iii) high antibiotic concentration ($\alpha_S(c) > \beta_S - \gamma x_S(t)$ for all $t \le \tau$). We now analyze these three parameter regimes separately:

(i) *Low antibiotic concentration ($\alpha_S(c) < \beta_S - \gamma x_S(t)$ for all $t \ge 0$):*

This situation is covered by a standard application of the technique reviewed in Section B.1. Precisely, with $\rho_k$ as introduced above and $s = \rho_R - \rho_S$, we have

$$\varphi^{\mathrm{b}}_{\mathrm{bs}}(\tau) = \max\left(0, \frac{s\rho_S\rho_R}{s\rho_S\rho_R + (\delta_R+\alpha_R)(x_S(0)\gamma\rho_R(1-e^{-s\tau})) + s(\rho_S - x_S(0)\gamma)(1-e^{-\rho_R\tau})}\right). \tag{B.17}$$

(ii) *Intermediate antibiotic concentration ($\alpha_S(c) > \beta_S - \gamma x_S(t)$ initially and then switches sign for some $T_S < \tau$):*

In this situation there are two critical times to consider. First, we need to identify the time $T_R$ that denotes the time from which on the growth rate of the resistant strain is positive, i.e.,

$$T_R = \min\left(\tau, \inf_{t\ge 0}\left\{x_S(0)e^{-\delta_S t} = (\beta_R - \alpha_R(c))\right\}\right), \tag{B.18}$$

where $x_S(0)e^{-\delta_S t} = x_S(t)$. The second time, denoted $T_S$, is the analogously defined time for the sensitive strain:

$$T_S = \min\left(\tau, \inf_{t\geq 0}\left\{x_S(0)e^{-\delta_S t} = (\beta_S - \alpha_S(c))\right\}\right). \tag{B.19}$$

Note that because of $\alpha_R < \alpha_S$ and because $\alpha_S > \beta_S - \gamma x_S(0)$, we always have $0 \leq T_R < T_S$, at least if the difference between $\beta_S$ and $\beta_R$ is not too large (which is never the case for our choice of parameters). The survival probability of the resistant strain until $T_R$ is given by the probability of the resistant cell not to die, i.e. $e^{-\delta_R T_R}$. In the time interval $[T_R, T_S]$ the resistant strain can start to grow and the sensitive strain still declines exponentially. To compute the probability of survival of the resistant strain in this time interval, one needs to solve Eq. (B.3) with integral boundaries $T_R$ and $T_S$. Unfortunately, this integral cannot be solved analytically. More details are provided in the next scenario (case (iii) - high antibiotic concentration). Here, we only state the probability of survival in this time interval, given by

$$\frac{1}{1 + \int_{T_R}^{T_S} \delta_R \exp\left(-\rho_R t + \gamma x_S(0)(1 - e^{-\delta_S(t+T_R)})/\delta_S\right) dt}. \tag{B.20}$$

Lastly, we need to combine this probability with the survival probability of the resistant strain in the time interval $[T_S, \tau]$. Since the growth rate of the resistant strain was positive in the interval $[T_R, T_S]$, the resistant strain can potentially have reproduced. To account analytically for all these possible trajectories, we make the approximation that from 10 resistant cells on, the resistant strain survives almost surely, i.e. with probability one, until the end of treatment. To compute the probability of the resistant strain to be composed of $j$ cells at time $T_S$, we make use of the probability generating function, defined by

$$G(z) = \mathbb{E}\left[e^{zX_R}\right], \tag{B.21}$$

where $X_R$ is the random variable denoting the number of resistant cells at time $T_S$. Then the probability of $X_R = j$ can be computed by

$$\mathbb{P}(X = k) = \left.\frac{\partial G^j(z)}{j!\partial^j z}\right|_{z=0}. \tag{B.22}$$

In our case, the derivatives simplify to (calculation in the *Mathematica* notebook 'derivatives_gen_funcs')

$$\mathbb{P}(X_R = j) = \frac{AB^{k-1}}{(A+B)^{k+1}}, \tag{B.23}$$

where the coefficients $A$ and $B$ are given by (we refer also to Uecker and Hermisson (2011), Eqs. (9)-(12))

$$\begin{aligned} A &= \exp\left(\frac{x_S(0)\gamma\left(1 - e^{-\delta_S(T_S - T_R)}\right)}{\delta_S} - \rho_R(T_S - T_R)\right), \\ B &= 1 - A + \int_{T_R}^{T_S} \delta_R \exp\left(-\rho_R t + \gamma x_S(0)(1 - e^{-\delta_S(t+T_R)})/\delta_S\right) dt. \end{aligned} \tag{B.24}$$

Note that the probability for no offspring, which basically means that the resistant strain dies out during $[T_R, T_S]$ is (Uecker and Hermisson (2011), Eq. (13))

$$\mathbb{P}(X_R = 0) = 1 - \frac{1}{A+B}. \tag{B.25}$$

Lastly, we still need to compute the probability for the resistant strain to survive during the time interval $[T_S, \tau]$. We again use one of the formulas that were derived in Uecker and Hermisson (2011) (Eq. (13)). For this, we denote the density-independent growth by $\rho_k = \beta_k - \alpha_k - \delta_k$ and the selective advantage of the resistant type by $s = \rho_R - \rho_S$. Then the extinction probability of a single resistant cell in the time interval $[T_S, \tau]$ is given by

$$\begin{aligned} p_{\text{ext}} &= \frac{\int_{T_S}^{\tau} \frac{\delta_R}{x_S(t)} e^{-st} dt}{1 + \int_{T_S}^{\tau} \frac{\delta_R}{x_S(t)} e^{-st} dt} \\ &= \delta_R \frac{\rho_R \gamma x_S(T_S)(1 - e^{-s(\tau - T_S)}) - s(\gamma x_S(T_S) - \rho_S)(1 - e^{-\rho_R(\tau - T_S)})}{\rho_R \rho_S s + \delta_R(\rho_R \gamma x_S(T_S)(1 - e^{-s(\tau - T_S)}) - s(\gamma x_S(T_S) - \rho_S)(1 - e^{-\rho_R(\tau - T_S)})}, \end{aligned} \tag{B.26}$$

where $x_S(t)$ is the deterministic trajectory of the density of the sensitive strain, here

$$x_S(t) = \frac{x_S(0)\rho_S e^{\rho_S t}}{\rho_S + \gamma x_S(0)(e^{\rho_S t} - 1)}. \tag{B.27}$$

The extinction probability when starting with $j$ resistant cells at time $T_S$ is given by $p_{\text{ext}}^j$.

Now, the overall survival probability of the resistant strain computes to

$$\varphi_{\text{bs}}^{\text{b}}(\tau) = \underbrace{e^{-\delta_R T_R}}_{t \le T_R} \left( \underbrace{\sum_{j=1}^{10} \mathbb{P}(X_R = j)(1 - p_{\text{ext}}^j)}_{\text{up to 10 offspring}} + \underbrace{\left(1 - \sum_{j=0}^{10} \mathbb{P}(X_R = j)\right)}_{\text{more than 10 offspring}} \right). \tag{B.28}$$

(iii) *High antibiotic concentration ($\alpha_S(c) > \beta_S - \gamma x_S(t)$ for all $t \le \tau$):*

In this situation, we need to distinguish between the scenario where (a) $\beta_R - \alpha_R(c) < 0$ and (b) $\beta_R - \alpha_R(c) > 0$.

In scenario (a), the growth rate of the resistant strain, $\lambda(t)$, is zero for all $t \ge 0$, no matter how small the sensitive population. The survival probability is then just given by resistant cell not dying until the end of treatment, i.e.,

$$\varphi_{\text{bs}}^{\text{b}}(\tau) = e^{-\delta_R \tau}, \tag{B.29}$$

which tends to zero for $\tau \to \infty$.

In case (b), the growth rate of the resistant strain will be positive for certain population sizes of the sensitive strain. Hence, it exists a $T_R \ge 0$, so that $\lambda(t) > 0$ for all $T_R > t$. This time $T$ is defined by

$$T_R = \min\left(\tau, \inf_{t \ge 0}\left\{x_S(0)e^{-\delta_S t} = (\beta_R - \alpha_R(c))\right\}\right), \tag{B.30}$$

where $x_S(0)e^{-\delta_S t} = x_S(t)$.

For times smaller than $T_R$, the growth rate of the resistant strain is zero. Therefore its survival probability until time $T_R$ is, as above, $e^{-\delta_R T_R}$. For the overall survival of the resistant strain until the end of treatment, we need to compute the survival of the resistant strain in the time interval $[T_R, \tau]$. For this, we need to solve the integral in Eq. (B.3). Unfortunately, this is not possible analytically because of the time dependency introduced by the term $\gamma x_S(t)$, which cannot be resolved in this case. The survival probability then reads

$$\varphi_{\text{bs}}^{\text{b}}(\tau) = \frac{e^{-\delta_R T_R}}{1 + \int_{T_R}^{\tau} \delta_R \exp\left(-\rho_R t + \gamma x_S(0)(1 - e^{-\delta_S(t + T_R)})/\delta_S\right) dt}. \tag{B.31}$$

##### B.3.2 Biocidal antibiotic

In this last case, the antibiotic drug increases the death rate (biocidal drug). Then the birth rate of a resistant cell is

$$\lambda_R(x_S(t), x_R(t)) = \max\left(\beta_R - \gamma_R x_S(t), 0\right) \tag{B.32}$$

and the death rate is

$$\mu_R(x_S(t), x_R(t)) = \delta_R + \alpha_R(c). \tag{B.33}$$

We can directly apply the general theory from Section B.1. The reason is that the sensitive strain is always sufficiently small so that $\lambda_R(t) > 0$ for all times $t$. In general, this condition being satisfied of course depends on the choice of the parameters $\beta_R, \gamma_R$ and the antibiotic-free steady state of $x_S$. In our parameter set of the simulations, we can ignore the maximum-condition in Eq. (B.32) because $\beta_R > \gamma_R x_S(t)$ for all $t \geq 0$. If this condition is not satisfied, we instead need to first compute the survival probability for the resistant strain to survive until its growth rate becomes positive, similar to the previous scenario. Once the growth rate $\lambda_R$ is positive, the survival analysis for the remaining time follows again along the lines in Section B.1.

Assuming again equal competition rates $\gamma_S = \gamma_R = \gamma$, the growth rates are given by $\rho_k = \beta_k - \alpha_k - \delta_k$ and the selective advantage of the resistant strain is $s = \max(0, \rho_R - \rho_S)$. Plugging these values into the general theory (Eq. (B.3)), we find for the survival probability until time $\tau$ of a resistant strain under a biocidal antibiotic

$$\varphi^{\mathrm{b}}_{\mathrm{bc}}(\tau) = \frac{s\rho_S\rho_R}{s\rho_R\rho_S + (\delta_R + \alpha_R)\left(x_S(0)\gamma\rho_R\left(1 - e^{-s\tau}\right) + s(\rho_S - x_S(0)\gamma)\left(1 - e^{-\rho_R\tau}\right)\right)}. \tag{B.34}$$

For $\tau \to \infty$, this survival probability simplifies to

$$\varphi^{\mathrm{b}}_{\mathrm{bc}}(\infty) = \frac{s\rho_S\rho_R}{s\rho_R\rho_S + (\delta_R + \alpha_R)\rho_S\left(x_S(0)\gamma + s\right)} = \frac{1}{1 + \frac{\delta_R + \alpha_R}{\rho_R}\left(\frac{x_S(0)\gamma}{s} + 1\right)}. \tag{B.35}$$

### C Predicting the antibiotic concentration minimizing resistant strain survival

Our goal is to derive Eq. (3) from the main text. This equality is a condition for the antibiotic concentration to maximize the survival probability of the resistant strain until the end of treatment. The general approach is as follows: First, we note that the general structure of the survival probability in all scenarios except biostatic drug with birth competition is of the form:

$$\varphi(c) = \frac{1}{1+f(c)}, \tag{C.1}$$

where $f(c)$ is a function that depends on the specific scenario. Compare this equation with Eqs. (B.9), (B.13) and (B.35) to find this structural form. The derivative of the survival probability with respect to the antibiotic concentration is then

$$\varphi'(c) = -\frac{f'(c)}{(1+f(c))^2}, \tag{C.2}$$

so that

$$\varphi'(c) = 0 \qquad \Leftrightarrow \qquad f'(c) = 0. \tag{C.3}$$

This is the equality, which we solve in the following, to obtain analytical conditions on the quantities involved.

#### C.1 Scenario: Biocidal drug, birth competition

Recall that $s(c) = \rho_R(c) - \rho_S(c)$ and that $\rho_k(c) = \beta_k - \delta_k - \alpha_k(c)$ for $k \in \{S, R\}$. Then we have

$$f(c) = \frac{(\alpha_R(c)+\delta_R)}{\rho_R(c)}\left(1+\frac{\gamma x_S(0)}{s(c)}\right). \tag{C.4}$$

Using $s'(c) = \alpha'_S(c) - \alpha'_R(c)$, we can write the derivative as (see also the *Mathematica* notebook 'concentrationLocation_survProb' to verify this equation)

$$f'(c) = \frac{x_S(0)\gamma\rho_R(c)(\delta_R+\alpha_R(c))(\alpha'_R(c)-\alpha'_S(c)) + \beta_R(\rho_R(c)-\rho_S(c))\alpha'_R(c)(x_S(0)\gamma+\rho_R(c)-\rho_S(c))}{(\rho_R(c)-\rho_S(c))^2\rho_R(c)^2}. \tag{C.5}$$

Solving for $f'(c) = 0$, we obtain

$$\alpha'_S(c) = \alpha'_R(c)\left(1+\frac{\beta_R(\rho_R(c)-\rho_S(c))}{\rho_R(c)(\delta_R+\alpha_R(c))}+\frac{\beta_R(\rho_R(c)-\rho_S(c))^2}{x_S(0)\gamma\rho_R(c)(\delta_R+\alpha_R(c))}\right). \tag{C.6}$$

Reordering the terms, we find Eq. (3) from the main text.

#### C.2 Scenario: Biocidal drug, death competition

In this scenario, the function $f(c)$ is given by

$$f(c) = \frac{x_S(0)\gamma(\delta_R+\alpha_R(c)+\rho_S(c))}{\rho_S(c)s(c)} - \frac{(\delta_R+\alpha_R(c))(\gamma x_S(0)-\rho_S(c))}{\rho_S(c)\rho_R(c)}. \tag{C.7}$$

The derivative is

$$f'(c) = \frac{1}{s(c)^2\rho_S(c)^2\rho_R(c)^2}\Big(\rho_S(c)\,\big[\rho_R(c)(x_S(0)\gamma\rho_R(c)(\delta_R+\alpha_R(c)+\rho_S(c))(\alpha'_R(c)-\alpha'_S(c)) \\ +(\rho_R(c)-\rho_S(c))\rho_S(c)(x_S(0)\gamma+(\rho_R(c)-\rho_S(c)))\alpha'_R(c)) \\ +(\rho_R(c)-\rho_S(c))^2(\delta_R+\alpha_R(c))(\rho_S(c)-x_S(0)\gamma)\alpha'_R(c)\big] \\ +x_S(0)\gamma(\rho_R(c)-\rho_S(c))(\delta_R+\alpha_R(c))\rho_R(c)\rho_S(c)\alpha'_S(c)\Big). \tag{C.8}$$

Solving for $f'(c) = 0$, we find

$$\alpha'_S(c) = \alpha'_R(c)\left(2 - \frac{(\rho_R(c)-\rho_S(c))^2 - x_S(0)\gamma\rho_S(c)}{x_S(0)\gamma\rho_R(c)}\right). \tag{C.9}$$

##### C.3 Scenario: Biostatic drug, death competition

This scenario is a bit more involved to analyze. Still, we start as before with the auxiliary function $f(c)$. Writing $s(c) = \beta_R - \delta_R - \alpha_R(c) - \beta_S + \delta_S + \alpha_S(c)$, we have

$$f(c) = \frac{x_S(0)\gamma(\delta_R+\rho_S(c))}{\rho_S(c)s(c)} - \frac{\delta_R(\gamma x_S(0)-\rho_S(c))}{\rho_S(c)\rho_R(c)}. \tag{C.10}$$

The derivative of this function is

$$f'(c) = \frac{\rho_S(c)}{s(c)^2\rho_S(c)^2\rho_R(c)^2}\left(\underbrace{x_S(0)\gamma\rho_R(c)^2(\delta_R+\rho_S(c))(\rho'_S(c)-\rho'_R(c))}_{(i)} \\ +\underbrace{\delta_R(\rho_R(c)-\rho_S(c))^2(x_S(0)\gamma-\rho_S(c))\rho'_R(c)}_{(ii)} \\ \underbrace{-x_S(0)\gamma\delta_R(\rho_R(c)-\rho_S(c))\rho_R(c)\rho'_S(c)}_{(iii)}\right). \tag{C.11}$$

It is not straightforward to solve $f'(c) = 0$ because of the maxima in the exponential growth rates $\rho_S$ and $\rho_R$, e.g.

$$\rho_S(c) = \max(0, \beta_S - \alpha_S(c)) - \delta_S. \tag{C.12}$$

Instead of finding a general solution to $f(c) = 0$, we derive a condition to locate the maximizing concentration.

We first define $\tilde{c}$ as the minimal concentration for which $\rho_S(c) = -\delta_S$, i.e., $\tilde{c} = \inf\{c : \rho_S(c) = -\delta_S\}$. Then for all concentrations above $\tilde{c}$, we have $\rho_S(c) = -\delta_S$ and hence $\rho'_S(c) = 0$ for $c > \tilde{c}$. We now argue that $f'(c) > 0$ for $c > \tilde{c}$, which implies that $\varphi^{\mathrm{b}}_{\mathrm{bs}}$ is decreasing (Eq. (C.2)) and thus attains its maximum at $c \le \tilde{c}$.

We study the terms $(i)-(iii)$ separately. First, we note that $\delta_R \le \delta_S = -\rho_S(c)$ because either $\delta_S = \delta_R$ or, assuming the cost of resistance being mediated by an increased death rate, then $\delta_R < \delta_S = -\rho_S$. We then have $(i) \le 0$ for $c > \tilde{c}$ because $\rho'_S(c) - \rho'_R(c) = -\rho'_R(c) > 0$ and $\delta_R + \rho_S(c) \le 0$. Second, we have $(ii) < 0$ because $\rho'_R(c) < 0$ and all other terms are positive. Lastly, we have $(iii) = 0$ because $\rho'_S(c) = 0$. Therefore, multiplication of the sum of $(i)-(iii)$ with $\rho_S(c) < 0$, we find that $f'(c) > 0$ for $c > \tilde{c}$. This implies that the maximizing concentration in this scenario (biostatic treatment and density affecting the death rate) will always be below $\tilde{c}$.

In our parameterization, we find that the maximizing concentration is exactly at $c = \tilde{c}$ because for $c < \tilde{c}$ the function $f'(c) < 0$, which can be evaluated numerically. If we assume that the death rate of the cells is equal to zero, then the maximizing concentration will be smaller or equal than the MIC of the sensitive strain, i.e., $\tilde{c} = \text{mic}_S$.

#### D Resistant population size at end of treatment

We now estimate the average population size of the resistant strain at the end of treatment, conditioned on its survival. We will perform all the computations on the density scale and in the end transform the density back to the numbers of bacterial cells, which we call the size. To do so, we simply multiply the computed quantities by the parameter $K$, which one can think of as the volume of the studied location.

To compute the density at the end of treatment, we replace the initial density of the resistant subpopulation by the *effective* initial density (Uecker and Hermisson, 2011). This value accounts for the conditioning of the trajectories on survival, which practically results in a larger population size than the deterministic prediction. The average of the effective initial density of the resistant subpopulation is given by $x_R(0)/\varphi(\tau)$, i.e. a rescaling by the survival probability. Formally, if we denote the effective initial population density by $v$, the distribution of this random variable is given by Eq. (40) from Uecker and Hermisson (2011):

$$\mathbb{P}(v \le v_0 | \text{ survival}) = 1 - \exp(-\varphi(\tau)v_0)\,, \tag{D.1}$$

which shows that the effective initial population size is exponentially distributed with parameter $\varphi(\tau)$. This is true if the resistant population size can initially be approximated by an exponentially growing population with rate $a(t) = \max(0, \max(0, \beta_R - \gamma x_S(t)) - \delta_R - \alpha_R(c))$ (in the case of biocidal treatment and density affecting the birth rate).

Since the trajectory is only approximated well by the exponential growth curve while the number of resistant cells is still rare, we compute the average time for the resistant subpopulation to reach a certain intermediate size. Here, we choose the threshold $x_R^c = 1/(K\varphi(\tau))$, which intuitively corresponds to the population size from which, on average, one cell (divided by $K$ to transform to a density) will found a surviving lineage. After hitting $x_R^c$, the trajectory follows the deterministic trajectory defined by the competitive Lotka-Volterra-like model described in Section A. It therefore remains to compute the mean of the hitting time of $x_R^c$, which we denote by $t_R^c$.

The distribution of $t_R^c$ can be computed from the distribution of the effective initial population density (Eq. (D.1)). Specifically, we have

$$\mathbb{P}(t_R^c \le t) = \mathbb{P}\left(a^{-1}\left(\ln\left(\frac{x_R^c}{v}\right)\right) \le t\right) = \mathbb{P}\left(v > x_R^c \exp(-a(t))\right) = \underbrace{\exp\left(-\varphi(\tau)x_R^c \exp(-a(t)\right)}_{=A(t)}. \tag{D.2}$$

The average hitting time is then given by

$$\mathbb{E}[t_R^c] = \int_0^\infty tA'(t)dt = \int_0^\infty t\exp\left(-\varphi(\tau)x_R^c\exp(-a(t))\right)\varphi(\tau)x_R^c\exp(-a(t))a'(t)dt. \tag{D.3}$$

The size of the resistant subpopulation is then computed as follows (scenario: biocidal treatment, density affecting birth rate):

(i) compute the mean hitting time $t_R^c$ (if $t_R^c > \tau$, set $t_R^c = \tau$);

(ii) set $\tilde{x}_R(0) = x_R^c$ and $\tilde{x}_S(0) = x_S(\mathbb{E}[t_R^c])$, where $x_S$ is given by the following differential equation:

$$dx_S/dt = x_S(\max(\beta_S - \gamma x_S, 0) - \alpha_S(c) - \delta_S)\,;$$

(iii) compute the value $\tilde{x}_R(\tau - \mathbb{E}[t_R^c])$, where $\tilde{x}_R$ and $\tilde{x}_S$ are given by the following dynamics

$$\frac{d\tilde{x}_S}{dt} = \tilde{x}_S\left(\max(\beta_S - \gamma(\tilde{x}_S + \tilde{x}_R), 0) - \alpha_S(c) - \delta_S\right),$$
$$\frac{d\tilde{x}_R}{dt} = \tilde{x}_R\left(\max(\beta_R - \gamma(\tilde{x}_S + \tilde{x}_R), 0) - \alpha_R(c) - \delta_R\right).$$

(iv) Multiply the resulting value by $K$ to translate the density into number of cells.

Because of the condition $t_R^c \leq \tau$, the theoretical curve will show discontinuities at the concentrations where this threshold is reached. These discontinuities are (unreasonable) jumps to higher values for the population size as shown in Fig. D.1. As visible, the discontinuities are not very pronounced when studying the size of the resistant subpopulation at the end of treatment, yet they substantially impact the carriage time of the resistant strain. To smoothen the carriage time curve at these discontinuities and by that improve the prediction, we also compute the trajectories of the resistant subpopulation if started at time 0 with the effective initial population size $x_R(0) = 1/(K\varphi(\tau))$. These population sizes are typically above the trajectories computed by steps (i)-(iii), except for the concentrations close to the discontinuities (compare dashed and dotted lines in Fig. D.1). To predict the size at the end of treatment in a reasonable way, we take the minimum of the values $\tilde{x}_R(\tau)$ (computed by (i)-(iii)) and $x_R(\tau)$ (computed by $x_R(0) = 1/(K\varphi(\tau))$), which smoothens the predictions a bit.

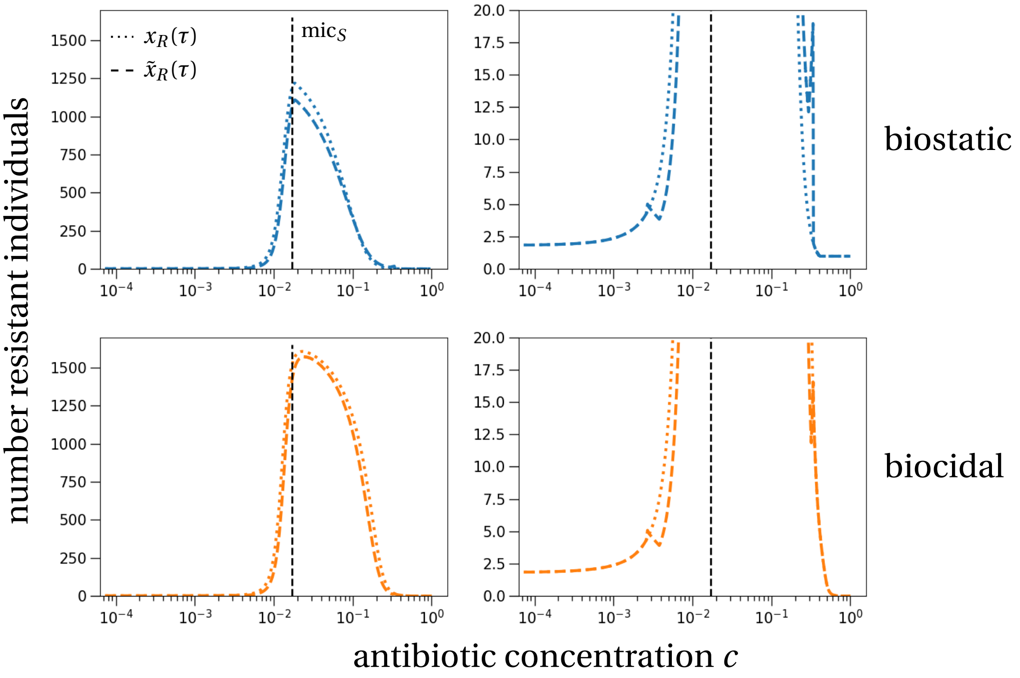


Figure D.1: **Different approximations for the resistant subpopulation size at the end of treatment.** The panels show approximations of the size at the end of treatment based on $x_R(\tau)$ (dotted lines) and the theoretically more accurate $\tilde{x}_R(\tau)$ (dashed lines). The dashed line shows discontinuities at the limits of the mutant selection window, which are not visible on the original scale (left vs. right column). By taking the minimal value of the two curves, we remain as accurate (theoretically) as possible for a large range of antibiotic concentrations, yet smoothen the prediction where the theory fails (upper bound of the mutant selection window). The vertical dashed line indicates the MIC of the sensitive strain. Parameters are our default parameter set as stated in Table A.2 with the MIC of the resistant strain being $\mathrm{mic}_R = 20 \times \mathrm{mic}_S$ and where density affects the birth rate.

#### D.1 Frequency of resistant strain at the end of treatment

In addition to the absolute size of the resistant subpopulation at the end of treatment, we also investigated the frequency of the resistant strain in the overall pathogen population at the end of treatment. The simulation results are shown in Fig. D.2.

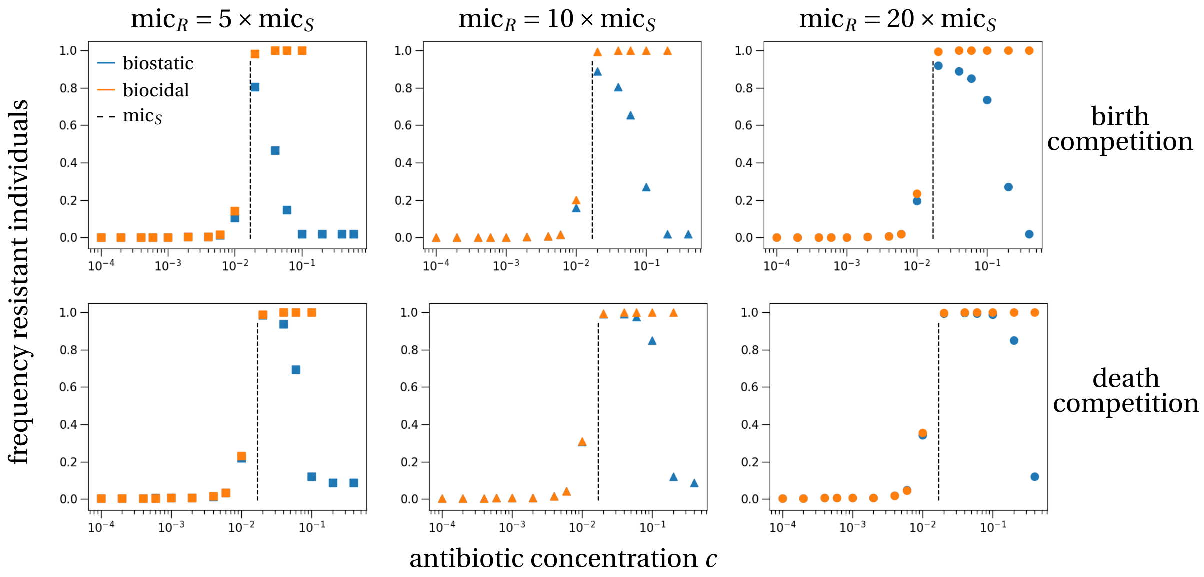


Figure D.2: **Resistant strain frequency at the end of treatment.** The frequency of resistant cells in the pathogen population shows a clear peak for biostatic treatment under variation of the antibiotic concentration (blue symbols). For biocidal antibiotic drugs (orange symbols) the frequency reaches the maximum close to the MIC of the sensitive strain and remains at a frequency of 1 for large antibiotic concentrations, i.e., no sensitive pathogens survive the treatment. Parameters are our default parameter set as stated in Table A.2.

### E Mean carriage time of the resistant population after treatment

If at the end of treatment a resistant subpopulation has survived and potentially even established, we ask how long it takes for the sensitive strain to replace the resistant strain in the host. To study this carriage time of the resistant strain, we use the framework of stochastic differential equations.

The equations of the densities of the sensitive and resistant strains, $x_S$ and $x_R$, that describe these dynamics are given by (for a derivation in this specific model we refer to Constable and McKane (2015), Huang et al. (2015) and Czuppon and Traulsen (2018)):

$$\begin{aligned} dx_S(t) &= x_S(t)(\lambda_S(x_S,x_R)-\mu_S(x_S,x_R))dt+\sqrt{\frac{x_S(t)(\lambda_S(x_S,x_R)+\mu_S(x_S,x_R))}{K}}dW_t^1, \\ dx_R(t) &= x_R(t)(\lambda_R(x_S,x_R)-\mu_R(x_S,x_R))dt+\sqrt{\frac{x_R(t)(\lambda_R(x_S,x_R)+\mu_R(x_S,x_R))}{K}}dW_t^2, \end{aligned} \tag{E.1}$$

where $\lambda_k(x_S,x_R)$ and $\mu_k(x_S,x_R)$ are the model-dependent birth and death rate as stated in Table A.1 without the effect of an antibiotic, and $W_t^1, W_t^2$ are independent Brownian motions. As in the previous sections, we work in the special case of equal competition rates, $\gamma_S=\gamma_R=\gamma$.

#### E.1 Reviewing the method of timescale separation in Lotka-Volterra systems

The basic idea of the method that we are going to use relies on a timescale separation where the fast timescale corresponds to the dynamics of the total population size ($x_S+x_R$), and the slow timescale corresponds to the frequency changes in the population close to its carrying capacity. The general theory has been formally proved by Katzenberger (1991) and recently been reviewed by Constable et al. (2013) and Parsons and Rogers (2017). The specific example of a Lotka-Volterra model has been analyzed in detail in Constable and McKane (2015). The fast population size changes, compared to slow relative frequency changes, stabilize the total population size very quickly so that the frequency dynamics are restricted to a center manifold that connects the two monotypic states, i.e., the states where either the resistant or the sensitive subpopulation has gone extinct. The dynamics on this center manifold can be obtained by projecting the frequency dynamics onto it, which results in a one dimensional stochastic differential equation. For one dimensional stochastic differential equations explicit solutions of fixation probabilities and mean extinction times exist (reviewed in Czuppon and Traulsen, 2021).

In the following, we apply the procedure as described in Constable and McKane (2015).

#### E.2 Death competition

We set $\rho_k=\beta_k-\delta_k$ with the index indicating the strain type, $k\in\{S,R\}$. We apply the following transformation: $y_S(t)=x_S(t)\,\gamma/\rho_S$, $y_R(t)=x_R(t)\,\gamma/\rho_R$ and rescale time by $\rho_S$ ($t\mapsto\rho_S t$). Eq. (E.1) then transforms to

$$\begin{aligned} dy_S(t) &= y_S(t)\left(1-y_S(t)-\frac{\rho_R}{\rho_S}y_R(t)\right)dt+\sqrt{\frac{\gamma(\beta_S+\delta_S+(\rho_S y_S(t)+\rho_R y_R(t)))y_S(t)}{\rho_S^2 K}}dW_t^1, \\ dy_R(t) &= y_R(t)\left(\frac{\rho_R}{\rho_S}-y_S(t)-\frac{\rho_R}{\rho_S}y_R(t)\right)dt+\sqrt{\frac{\gamma(\beta_R+\delta_R+(\rho_S y_S(t)+\rho_R y_R(t)))y_R(t)}{\rho_S\rho_R K}}dW_t^2. \end{aligned} \tag{E.2}$$

To apply the results on the mean extinction time of the resistant strain obtained in Constable and McKane (2015), we translate their parameters (their Eq. (16)) to our situation – their Greek letters correspond to our Latin letters and vice versa. In their notation, we have $b_S = d_S = c_{SS} = c_{RS} = 0$ and

$$b_R = \frac{\beta_R - \beta_S}{\varepsilon \beta_S}, \quad d_R = \frac{\delta_R - \delta_S}{\varepsilon \delta_S}, \quad c_{SR} = c_{RR} = \frac{\rho_R - \rho_S}{\varepsilon \rho_S}, \tag{E.3}$$

where $\varepsilon$ measures the fitness difference between the sensitive and the resistant strain. Note that with this choice of parameters, Eq. (17) in Constable and McKane (2015) is automatically satisfied, so that the boundary conditions, which correspond to the monotypic equilibria, are as required for the theoretical analysis to work. The results in Constable and McKane (2015) are exact for infinitely small fitness differences ($\varepsilon \to 0$), but are still a reasonably good approximation for fitness differences up to 10% (Czuppon and Traulsen, 2018).

Translating the results from Eqs. (14), (15) and (19) in Constable and McKane (2015) to our parameters, we obtain the following one-dimensional stochastic diffusion for the frequency of sensitive alleles in the population, $p = y_S$:

$$dp(t) = \left(1 - \frac{\rho_R}{\rho_S}\right) p(t)(1-p(t))dt + \sqrt{\frac{2\beta_S \gamma p(t)(1-p(t))}{\rho_S^2 K}} dW_t. \tag{E.4}$$

Note that the variable $y_S$ indeed corresponds to the frequency of the sensitive type because on the center manifold we have $y_S + y_R = 1$.

From this one dimensional stochastic differential equation, we can directly compute the mean time to extinction of the resistant allele (conditioned on extinction of the resistant strain). For this, we first need to compute the fixation probability of the sensitive strain when started with frequency $p$, denoted $\xi_{\text{fix}}(p)$, which is given by (e.g. Otto and Day, 2007; Czuppon and Traulsen, 2021)

$$\xi_{\text{fix}}(p) = \min\left(1, \frac{S(p) - S(0)}{S(1) - S(0)}\right), \tag{E.5}$$

where $S(p)$ is the scale function of the stochastic process corresponding to the stochastic diffusion in Eq. (E.4), given by

$$S(p) = \int^p \exp\left(-2\int^q \frac{\left(1-\frac{\rho_R}{\rho_S}\right) q'(1-q')}{\frac{2\beta_S\gamma q'(1-q')}{\rho_S^2 K}} dq'\right) = -\frac{\beta_S \gamma}{\rho_S K(\rho_S - \rho_R)} \exp\left(-\frac{\rho_S K(\rho_S - \rho_R)}{\beta_S \gamma} p\right). \tag{E.6}$$

Setting $C = \frac{\rho_S K(\rho_S - \rho_R)}{\beta_S \gamma}$, the fixation probability then simplifies to

$$\xi_{\text{fix}}(p) = \min\left(1, \frac{1 - \exp(-Cp)}{1 - \exp(-C)}\right). \tag{E.7}$$

We can now compute the mean extinction time of the resistant strain conditioned on its extinction, $T_{\text{ext}}$. We denote the infinitesimal variance in Eq. (E.4) by $\sigma^2(p) = \frac{2\beta_S\gamma p(1-p)}{\rho_S^2 K}$. If the initial frequency of the sensitive strain is $p_0$, the frequency at the end of the antibiotic treatment, we find (Otto and Day, 2007; Ewens, 2004)

$$T_{\text{ext}} = 2\int_0^{p_0} (1 - \xi_{\text{fix}}(p_0)) \frac{S(p) - S(0)}{\sigma^2(p) S'(p)} \frac{\xi_{\text{fix}}(p)}{\xi_{\text{fix}}(p_0)} dp + 2\int_{p_0}^1 \xi_{\text{fix}}(p_0) \frac{S(1) - S(p)}{\sigma^2(p) S'(p)} \frac{\xi_{\text{fix}}(p)}{\xi_{\text{fix}}(p_0)} dp. \tag{E.8}$$

Lastly, we need to account for the time rescaling that we used to derive Eq. (E.2), i.e., we divide $T_{\text{ext}}$ by $\rho_S$, which then gives the mean extinction time of the resistant strain conditioned on its extinction.

#### E.3 Birth competition

If the bacterial density affects the birth rates in the individual-based model (Table 1 in the main text), the stochastic differential equations describing the dynamics of the (transformed) sensitive and resistant subpopulations, $y_S$ and $y_R$, are

$$\begin{aligned} dy_S(t) &= y_S(t)\left(1 - y_S(t) - \frac{\rho_R}{\rho_S} y_R(t)\right)dt + \sqrt{\frac{\gamma(\beta_S + \delta_S - (\rho_S y_S(t) + \rho_R y_R(t)))y_S(t)}{\rho_S^2 K}}\, dW_t^1, \\ dy_R(t) &= y_R(t)\left(\frac{\rho_R}{\rho_S} - y_S(t) - \frac{\rho_R}{\rho_S} y_R(t)\right)dt + \sqrt{\frac{\gamma(\beta_R + \delta_R - (\rho_S y_S(t) + \rho_R y_R(t)))y_R(t)}{\rho_S \rho_R K}}\, dW_t^2. \end{aligned} \tag{E.9}$$

Note that the only difference to Eq. (E.2) is a minus sign in the variance term. This change in sign results in a different variance term in Eq. (E.4). Applying the approach from Constable and McKane (2015), specifically their Eqs. (7),(12) and (15), results in the following stochastic differential equation on the center manifold:

$$dp(t) = \left(1 - \frac{\rho_R}{\rho_S}\right)p(t)(1-p(t))dt + \sqrt{\frac{2\delta_S \gamma p(t)(1-p(t))}{\rho_S^2 K}}\, dW_t. \tag{E.10}$$

Consequently, the probability of fixation of the sensitive strain and the mean extinction time of the resistant strain change slightly. In detail, the scale function changes to

$$S(p) = -\frac{1}{C}\exp\left(-Cp\right), \quad \text{with} \quad C = \frac{\rho_S K(\rho_S - \rho_R)}{\delta_S \gamma}. \tag{E.11}$$

From here on, the analysis is analogous to the previous section, i.e., from Eq. (E.7) onwards.

#### E.4 Comparison between deterministic and stochastic prediction

We compare our stochastic prediction to carriage time estimates that are derived from the deterministic model. There are two (partly) deterministic estimates that we can consider: (i) a purely deterministic estimate, where both the treatment and post-treatment phase are modeled deterministically; (ii) a hybrid estimate, where the treatment phase is modeled stochastically, and the post-treatment phase deterministically. For the purely deterministic estimate (dotted lines in Fig. E.1), we observe that the carriage time is only defined in the mutant selection window because otherwise the resistant strain does not survive treatment. In contrast, for the hybrid prediction (dashed lines in Fig. E.1) we use the stochastic estimate of the resistant subpopulation at the end of treatment and then propagate the population dynamics deterministically. Hence, the initial condition of the pathogen population at the end of treatment is the same in the hybrid and the purely stochastic estimate. The discontinuity in the curves is explained by the discontinuous estimation of the subpopoulation size at the end of treatment, see the discussion at the end of Section D.

We generally observe that the deterministic predictions overestimate the carriage time for models where densitiy affects the death rate. In models where density affects the birth rate, the overestimation only becomes visible when the growth rate difference between the resistant and sensitive strain become small, e.g. in Fig. E.2 about 4%.

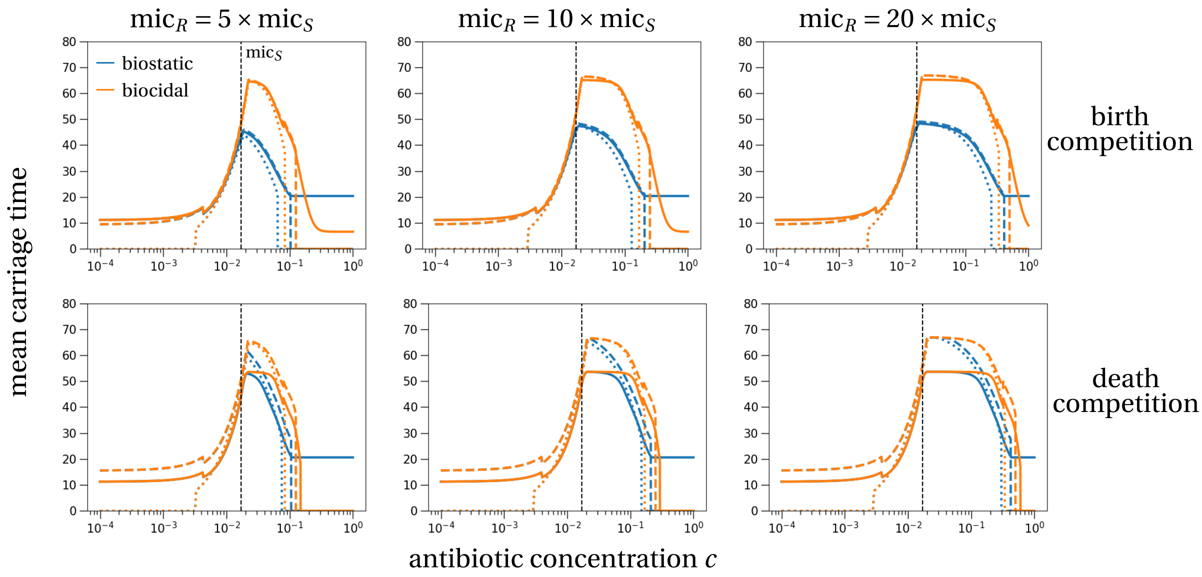


Figure E.1: **Comparison of deterministic and stochastic predictions of the carriage time.** The solid line corresponds to the purely stochastic prediction as used in the main text; the dashed line depicts the hybrid prediction, where the treatment phase is described stochastically and the post-treatment phase deterministically; the dotted line shows the purely deterministic prediction. Parameters are chosen according to our default parameter set as stated in Table A.2.

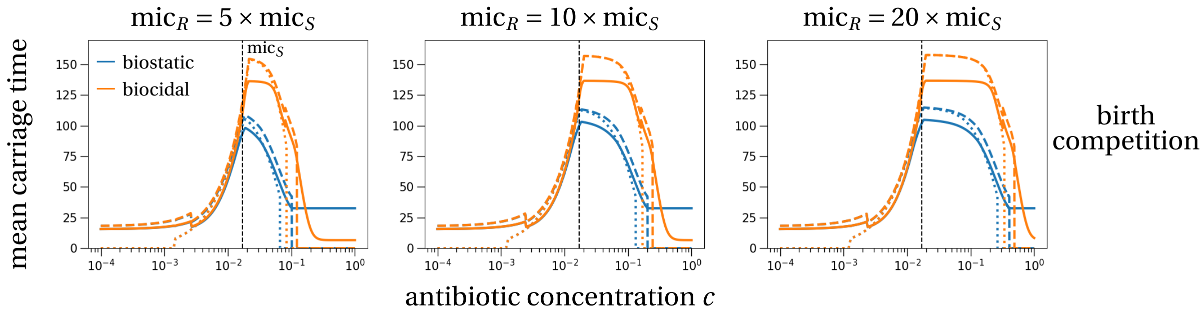


Figure E.2: **Comparison of deterministic and stochastic predictions of the carriage time for small growth rate differences between resistant and sensitive strain.** The same as Fig. E.1 but with $\beta_R = 2.4$.

### F De-novo emergence of antibiotic resistance

In this section, we study the *de novo* emergence of antibiotic resistance. Before, we have always assumed that at the beginning of treatment, a resistant cell was already present in the population, a situation referred to as standing genetic variation. Here, we investigate whether our main conclusions from the main text still hold in the situation of *de novo* emergence of resistance. In particular, we show that the maximal risk of emergence and highest resistant subpopulation size are still located close to the MIC of the sensitive strain (Figs. F.1-F.6), which indicates that our findings hold for all types of resistance emergence during treatment.

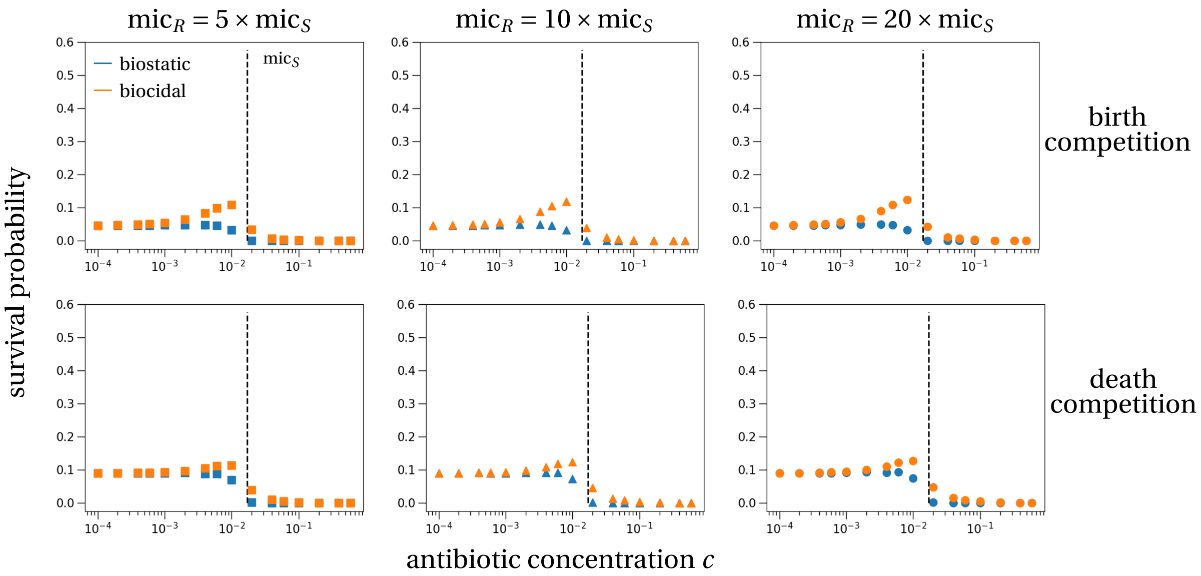


Figure F.1: **Survival probability of the resistant strain until the end of treatment.** In contrast to Fig. 3 in the main text, here initially there is no resistant cell in the population. Instead, at each birth event of a sensitive cell the offspring gains resistance with a certain probability reflecting mutations. The parameters are as stated in Table A.2 with the mutation probability set to $\eta = 1/(50K)$.

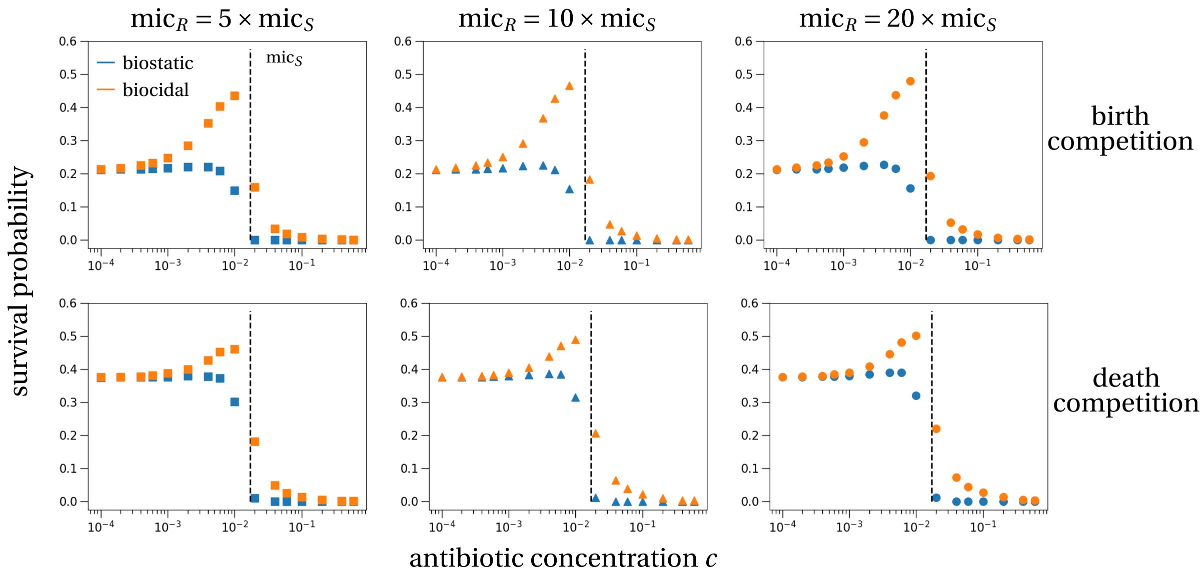


Figure F.2: **Survival probability of the resistant strain until the end of treatment.** The same as Fig. F.1, but with a mutation probability of $\eta = 1/(10K)$.

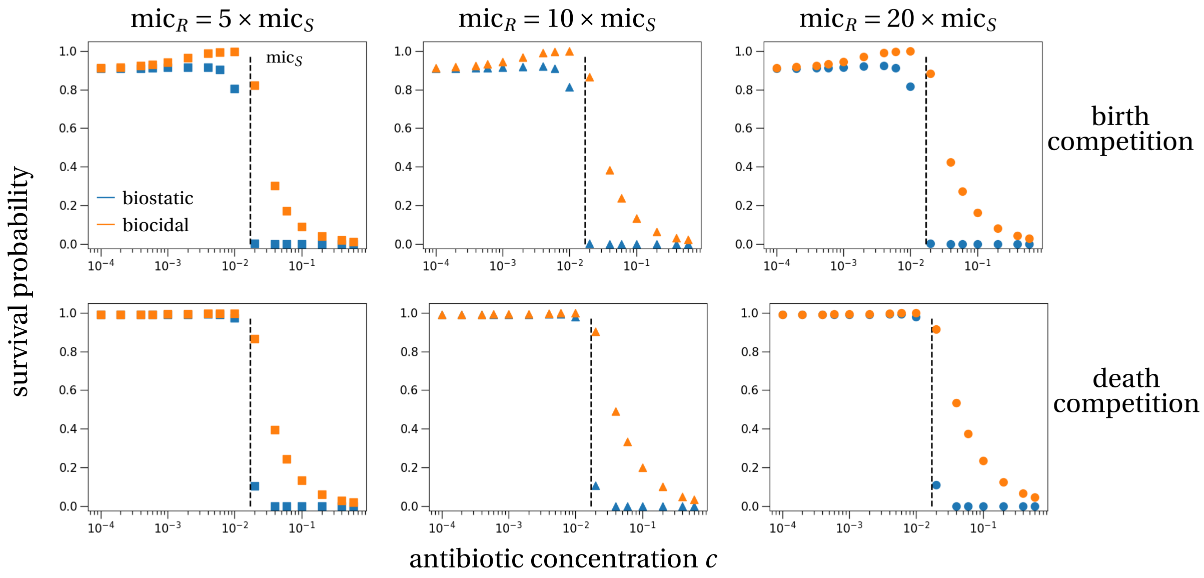


Figure F.3: **Survival probability of the resistant strain until the end of treatment.** The same as Fig. F.1, but with a mutation probability of $\eta = 1/K$.

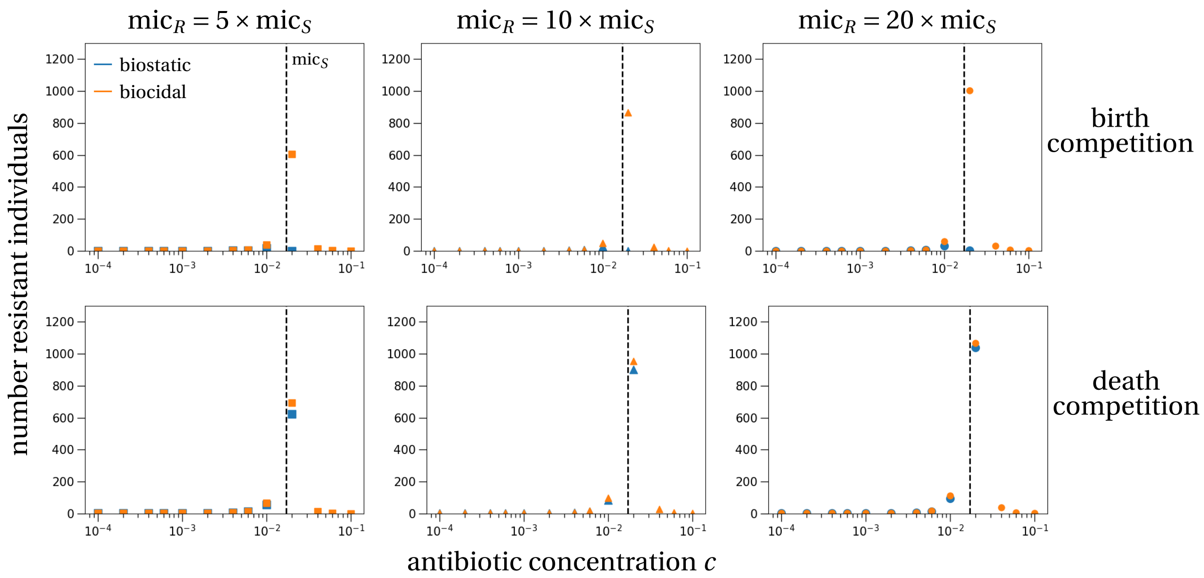


Figure F.4: **Size of the resistant subpopulation at the end of treatment.** In contrast to Fig. 4 in the main text, here initially there is no resistant cell in the population. Instead, at each birth event of a sensitive cell the offspring gains resistance with a certain probability reflecting mutations. The parameters are as stated in Table A.2 with the mutation probability set to $\eta = 1/(50K)$.

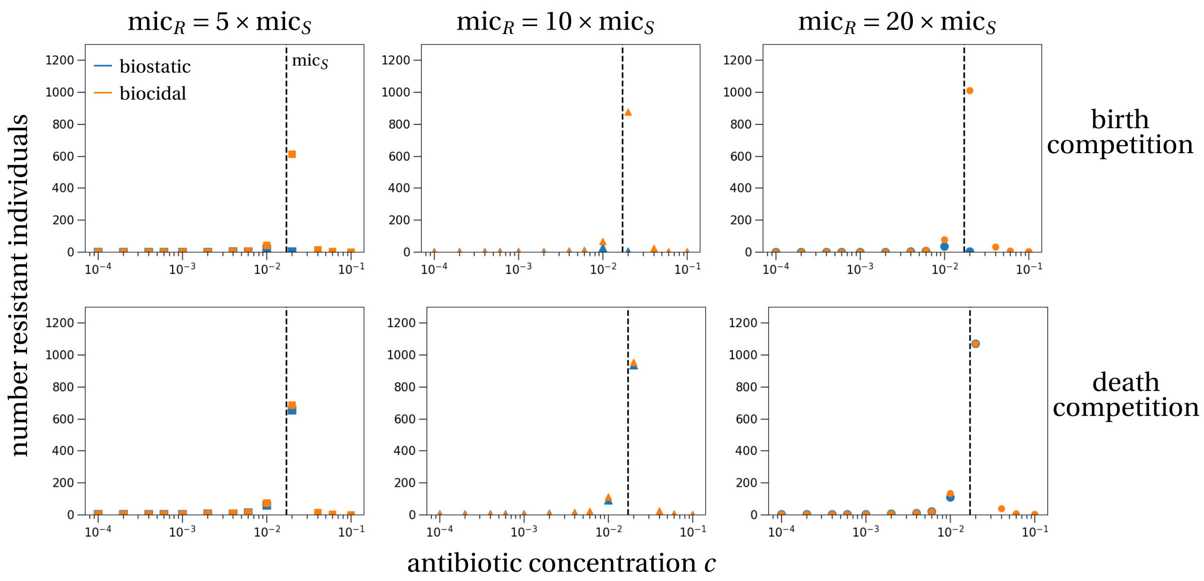


Figure F.5: **Size of the resistant subpopulation at the end of treatment.** The same as Fig. F.4, but with a mutation probability of $\eta = 1/(10K)$.

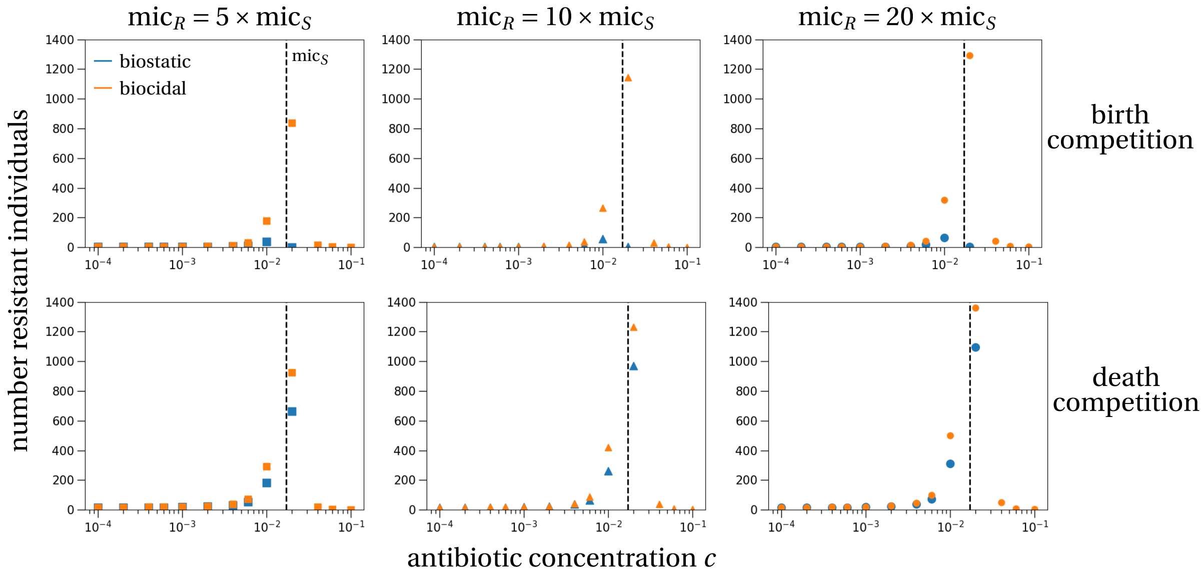


Figure F.6: **Size of the resistant subpopulation at the end of treatment.** The same as Fig. F.4, but with a mutation probability of $\eta = 1/K$.

### G Alternative parameterization of the model

In this section, we explore a parameterization that is more realistic for microbial bacteria. For the interpretation of our model, this means that these bacteria are under bystander selection, i.e., treatment is not administered to reduce this bacterial population.

The population dynamical parameters of the sensitive growth rate are set to reflect growth of for example *Escherichia coli* in the gut: growth rate = $\beta_S = 11$ per day, which corresponds to a replication time of 90 minutes. The death rate is set to $\delta_S = 0.5$, which reflects the outflux of the gut, on average two days (Poulsen et al., 1995). The growth rate of the resistant strain is set to $\beta_R = 10.6$, which corresponds to an approximately 10% fitness cost compared to the sensitive strain (Melnyk et al., 2015). In accordance with the methodology from Melnyk et al. (2015), we calculated the fitness cost by the relative growth of the resistant strain compared to the sensitive strain within 6 hours, i.e., $\exp(\beta_R \times 0.25) / \exp(\beta_S \times 0.25)$. The death rate of the resistant strain is assumed to be equal to the sensitive, $\delta_R = \delta_S = 0.5$.

The remaining parameters, in particular the antibiotic response curve, remain unchanged to our main scenario (Table A.2).

#### G.1 Survival probability

We first simulate and predict the survival probabilities for this alternative parameter set (Fig. G.1).

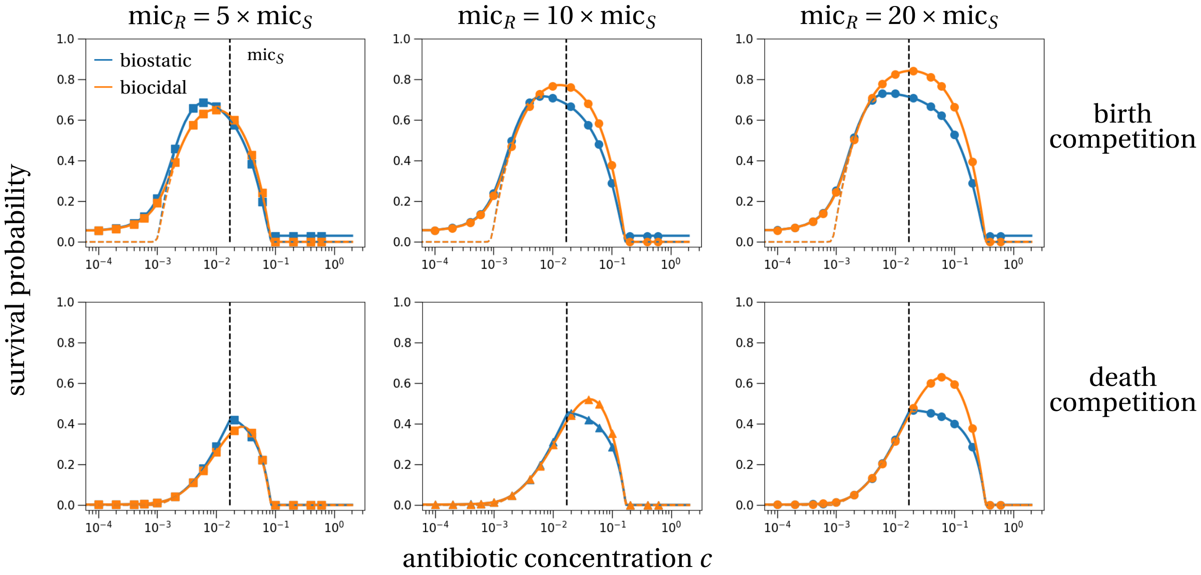


Figure G.1: **Survival probability of the resistant subpopulation for varying drug concentrations, MIC values, drug types and model assumptions.** At the beginning of treatment the population consists of the sensitive strain at its carrying capacity and a single resistant cell. Then treatment, either with a biostatic (blue) or biocidal (orange) antibiotic, is applied for seven days (solid lines) or infinitely long (colored dashed lines). The vertical dashed line indicates the MIC of the sensitive strain. Symbols show averages from $10^6$ stochastic simulations. Parameters correspond to the alternative parameterization with $\beta_S = 11$ ($\text{d}^{-1}$), $\beta_R = 10.6$ ($\text{d}^{-1}$), $\delta_S = \delta_R = 0.5$ ($\text{d}^{-1}$), and all remaining parameters as stated in Table A.2.

We observe a very similar pattern compared to the parameter set from the main text (Fig. 3 in the main text). In particular, birth competition results in higher survival probabilities of the resistant strain than death competition (compare upper and lower rows). Biostatic antibiotics suppress the resistant survival probability stronger than biocidal drugs, except for relatively small increases of resistance (left column) and concentrations below the sensitive MIC, which is consistent with Fig. 3 in the main text.

#### G.2 Size at the end of treatment

Next, we simulate and predict the size of the resistant subpopulation at the end of treatment, if it survives.

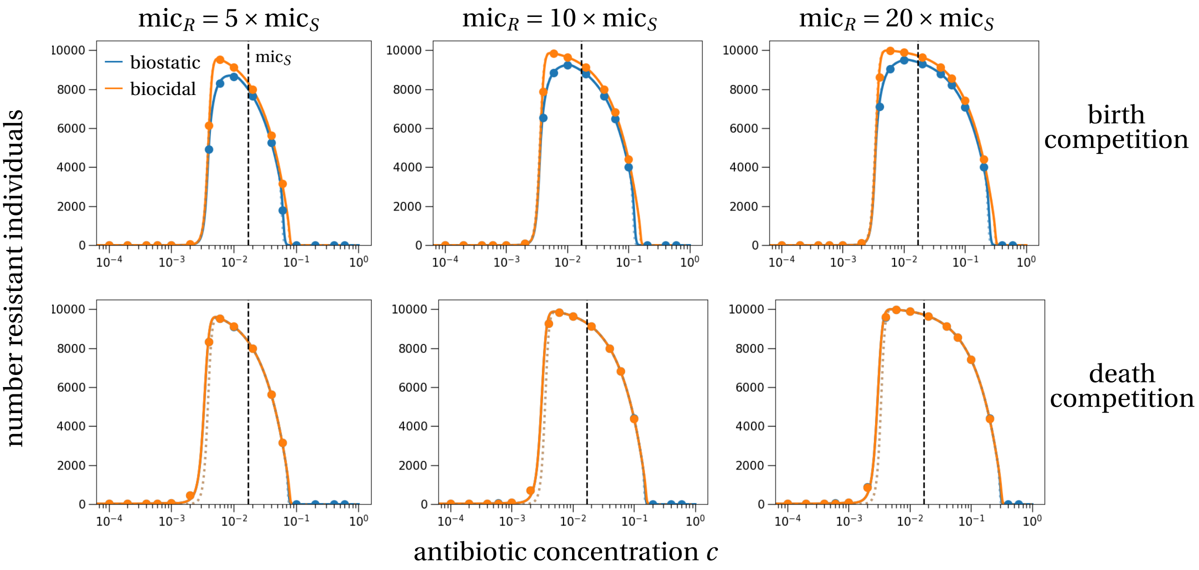


Figure G.2: **Size of the resistant subpopulation at the end of treatment** if the resistant subpopulation survives. At the onset of treatment there is exactly one resistant cell in the population and treatment lasts for seven days. The top row shows the results for the model of birth competition, the lower row corresponds to death competition. Dotted lines show the deterministic prediction of the resistant subpopulation size, solid lines correspond to the stochastic prediction that incorporates a 'correction' due to conditioning on survival. Symbols are the mean resistant subpopulation sizes of $10^6$ stochastic simulations that were conditioned on survival of the resistant subpopulation. Blue and orange colors correspond to biostatic and biocidal treatment, respectively. Parameters are as in Fig. G.1.

We find a similar qualitative pattern as in the default parameter set in Fig. 4 in the main text, quantitatively the resistant population sizes are strongly increased because of the higher resistant birth rate $\beta_R$. The only difference to the default parameter set is that the highest resistant population sizes are reached clearly below the sensitive MIC for all scenarios. As in the default parameter set, differences between biostatic and biocidal drugs are only visible for the model of birth competition and are negligible in the model of death competition.

#### G.3 Carriage time of the resistant strain

Next, we simulate and predict the carriage time of the resistant subpopulation after treatment, if it survives.

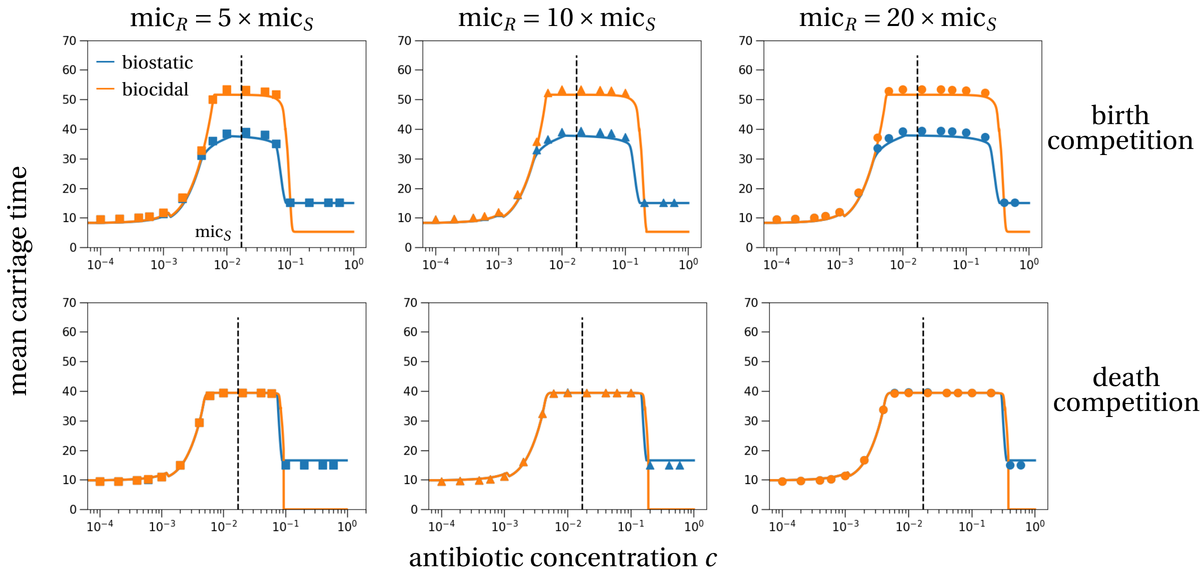


Figure G.3: **Mean carriage time of the resistant subpopulation in a host** (including the treatment time $\tau = 7$). The carriage time is set to zero if the resistant subpopulation did not survive antibiotic treatment. Symbols show the average carriage time of $10^6$ stochastic simulations. The color coding and figure structure is the same as in the previous figures. Parameters are as in Fig. G.1.

Compared to our main parameter set (Fig. 5 in the main text), we find reduced carriage times. This may be explained by the larger birth rates in this alternative parameter set, $\beta_S = 11$ instead of $\beta_S = 2.5$ in the main text, which results in a higher turnover of cells overall. This implies that the population size of the resistant subpopulation at the end of treatment is less good of a predictor about carriage times than the overall turnover of cells, which is defined as the sum of growth and death rates, $\lambda_S + \mu_S$. Besides this reduced carriage time, the main conclusions carry over to this alternative parameter set: biostatic drugs have a lower carriage time than bioicidal drugs under birth competition. This effect vanishes in the model of death competition. Lastly, the carriage time shows a plateau at concentrations above the size-maximizing concentration (compare to Fig. G.2).

#### H Alternative interaction of antibiotic and density-dependent processes

We study an alternative model for within-host population dynamics of the bacterial population, which has been used to study differences between biostatic and biocidal drugs under periodic antibiotic treatment (Marrec and Bitbol, 2020). In this model the antibiotic is interacting multiplicatively with the density-dependent regulation term, instead of additively as studied in the main text and in Section A above. For example, the sensitive strain dynamics in the case of biostatic treatment and birth competition are then given by

$$\frac{dx_S(t)}{dt} = x_S(t)\left(\max\left(0, (\tilde{\beta}_S - \alpha_S(c))\left(1-\tilde{\gamma}_S(x_S(t)+x_R(t))\right)\right) - \tilde{\delta}_S\right). \tag{H.1}$$

The birth and death rates in this model for all different scenarios are stated in Table H.1. Compared to the previous scenarios described in Section A, there are two new scenarios: biostatic treatment with birth competition, and biocidal treatment with death competition. The other two scenarios in Table H.1 are equivalent to previously studied scenarios, which is made clear by a parameter transformation. For example, in the scenario of biocidal treatment and birth competition, we can set $\tilde{\beta}_S = \beta_S$, $\tilde{\delta}_S = \delta_S$ and $\tilde{\gamma}_S = \gamma_S/\beta_S$ to recover the scenario in Section A.

We now study how the two new scenarios affect the population dynamics of the sensitive strain and the survival probability. Overall, our conclusions about the location antibiotic concentration, which maximizes the survival probability, being close to the MIC of the sensitive strain also hold under this alternative model.

##### H.1 Population dynamics

Compared to Section A the new curves for the sensitive subpopulation are the scenario of biostatic treatment and birth rate-affecting density regulation (solid blue lines in Fig. H.1) and the scenario of biocidal treatment and death rate-affecting density regulation (dotted orange lines). The former scenario results in a less strong effect of the antibiotic, compared to the original model, and the latter results in a stronger effect of the antibiotic on the population size (compare Figs. A.1 and H.1). These effects are explained by comparison of the respective birth and death rates in the different models. Investigating these inequalities yields a surprising result in this model: There is a positive interaction between the antibiotic and the density regulatory term, which results in an increased birth rate compared to the original model:

$$(\beta_S - \alpha_S)\left(1 - \frac{\gamma_S}{\beta_S}x_S\right) > \beta_S - \alpha_S - \gamma_S x_S \qquad \Leftrightarrow \qquad \alpha_S \frac{\gamma_S}{\beta_S} x_S \quad > 0. \tag{H.2}$$

| | | birth competition | death competition |
|---|---|---|---|
| **biostatic** | $\lambda_S(t)$ | $\max\left((\beta_S - \alpha_S(c))\left(1 - \frac{\gamma_S x_S(t)}{K}\right), 0\right)$ | $\max\left(\beta_S - \alpha_S(c), 0\right)$ |
| | $\mu_S(t)$ | $\delta_S$ | $\delta_S\left(1 + \frac{\gamma x_S(t)}{K}\right)$ |
| **biocidal** | $\lambda_S(t)$ | $\max\left(\beta_S\left(1 - \frac{\gamma_S x_S(t)}{K}\right), 0\right)$ | $\beta_S$ |
| | $\mu_S(t)$ | $\delta_S + \alpha_S(c)$ | $(\delta_S + \alpha_S(c))\left(1 + \frac{\gamma x_S(t)}{K}\right)$ |

Table H.1: **Alternative birth and death rates in the four studied scenarios.** This model has been studied in Marrec and Bitbol (2020).

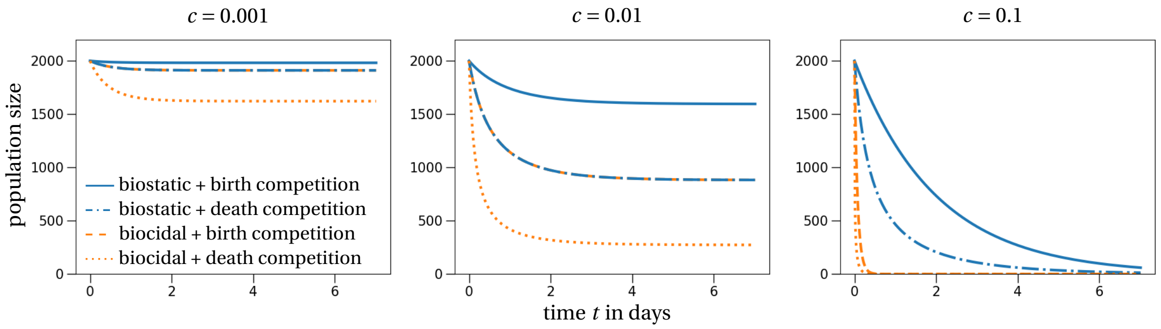


Figure H.1: **Population dynamics of sensitive subpopulation.** The population dynamics of the sensitive subpopulation are plotted for three different concentrations of the antibiotic, increasing from left to right. The birth and death rates of the different scenarios are given in Table H.1. The parameter values are as stated in Table A.2, except for the competition parameter $\gamma$. For biostatic treatment (blue lines), we set $\tilde{\gamma} = \gamma/\beta_S$, and for biocidal treatment (orange lines) $\tilde{\gamma} = \gamma/\delta_S$ to ensure comparable steady states of the sensitive subpopulation between this model and the model studied in the main text.

#### H.2 Survival probability

We simulate the survival probabilities in all the different scenarios as stated in Table H.1. We find no substantial difference between the alternative model implementation here and the original model from the main text (compare Fig. 3 with Fig. H.2). Of course, two scenarios (biostatic plus death competition and biocidal plus birth competition; blue symbols in bottom row and orange symbols in top row in Fig. H.2) are the same as in the original model, so no difference would be expected for these data points. The other two scenarios (biostatic plus birth competition and biocidal plus death competition; blue symbols in top row and orange symbols in bottom row in Fig. H.2) show differences in the survival probabilities, however not in the overall shape of the curve. Importantly, our conclusions about the location of the maximal survival probability being at concentrations close to the MIC of the sensitive strain translate to this model.

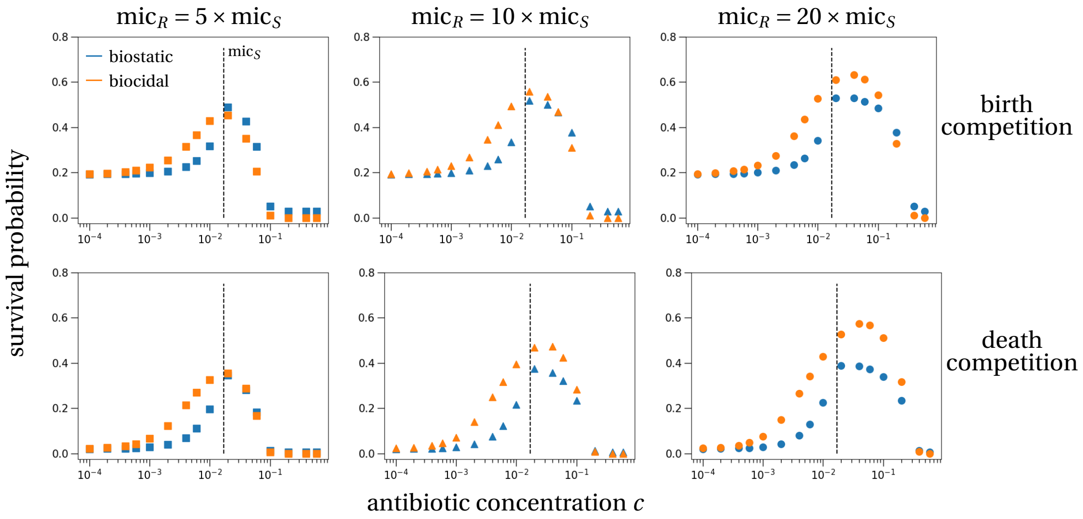


Figure H.2: **Survival probability of the resistant subpopulation in an alternative model.** The parameters are chosen according to the default parameter set (Table A.2) with the exception of the competition rate $\gamma$: top row $\tilde{\gamma} = \gamma/\beta_S$, bottom row $\tilde{\gamma} = \gamma/\delta_S$.

### I Alternative antibiotic response curve parameterizations

In this section we study different parameterizations of the antibiotic response curve. In the main text, we have used a parameterization that was motivated by the estimated parameter set for cyprofloxacin from the study by Regoes et al. (2004). Here, we maintain the same functional form for the antibiotic response curve, i.e.

$$\alpha_j(c) = (\psi_{j,\max} - \psi_{j,\min}) \frac{\left(\frac{c}{\mathrm{mic}_j}\right)^\kappa}{\left(\frac{c}{\mathrm{mic}_j}\right)^\kappa - \frac{\psi_{j,\min}}{\psi_{j,\max}}}, \tag{I.1}$$

but use different parameter sets to study if our conclusion about the location of the intermediate maximum are robust with respect to variations of the antibiotic response curve. In particular, we use the parameter sets for the antibiotics ampicillin and tetracycline, again from the study by Regoes et al. (2004), and additionally study a very steep and a very flat antibiotic response curve (parameters are stated in Table I.1). These curves are shown in Fig. I.1.

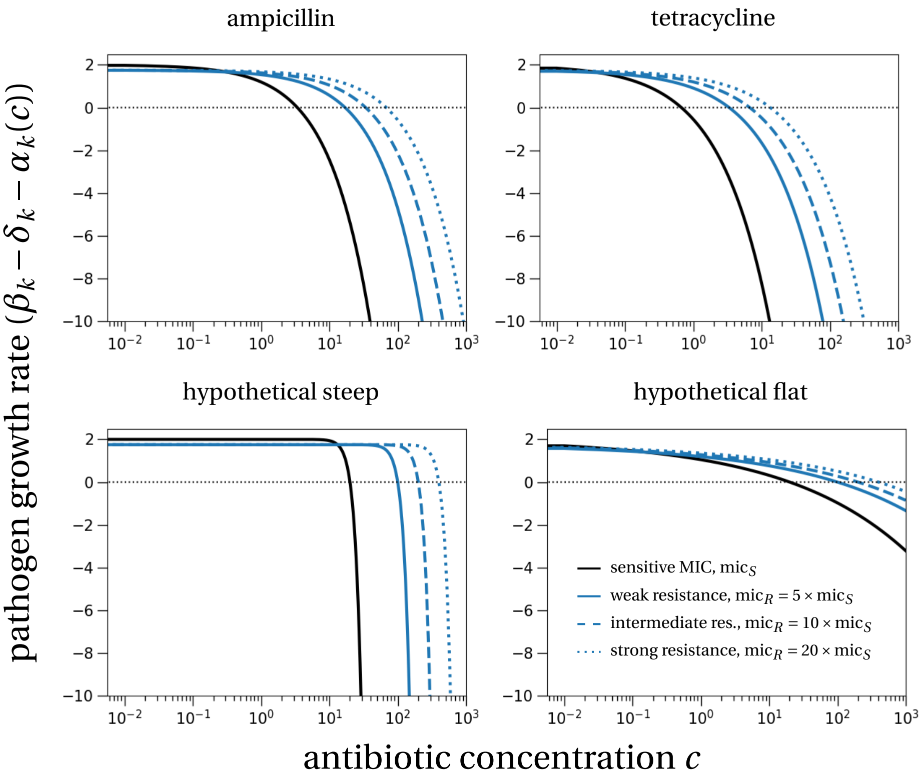


Figure I.1: **Different parameterizations of the antibiotic response curves.** The parameters are as stated in Table A.2, except for the parameters defining the antibiotic response $\alpha(c)$, which are stated in Table I.1.

#### I.1 Survival probability

The survival probability for the different antibiotic response parameterizations does not vary substantially from the findings in the main text (Fig. 3). The general pattern seems to be that the steeper the antibiotic response curve, i.e., the larger $\kappa$, the more shifted to the right, compared to the sensitive MIC, are the curves showing the probability of survival (Figs. I.2-I.5).

| **Parameter** | ampicillin | tetracycline | Hypothetical steep | Hypothetical flat |
|---|---|---|---|---|
| $\mathrm{mic}_S$ | 3.4 | 0.67 | 20 | 20 |
| $\psi_{\min}$ | $-96\times\log(10)$ | $-194\times\log(10)$ | $-72\times\log(10)$ | $-72\times\log(10)$ |
| $\kappa$ | 0.75 | 0.61 | 5 | 0.25 |

Table I.1: **Alternative parameterizations of the antibiotic response curve.**

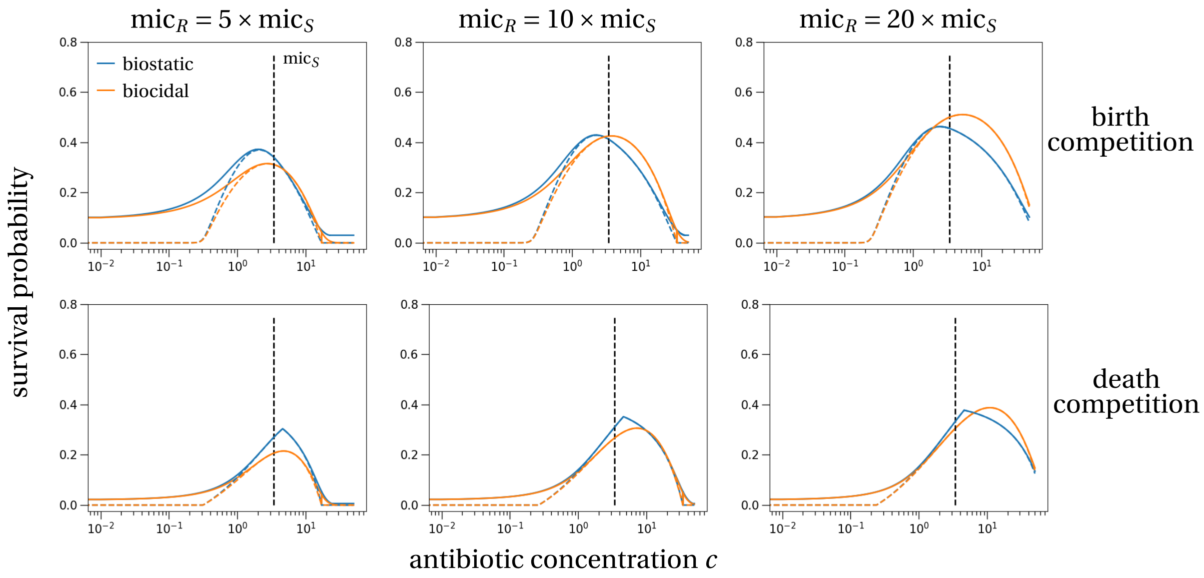


Figure I.2: **Survival probability for the parameterization with ampicillin.** The solid line shows the theoretical prediction for a treatment of seven days, the dashed lines show the survival probabilities for an infinitely long treatment.

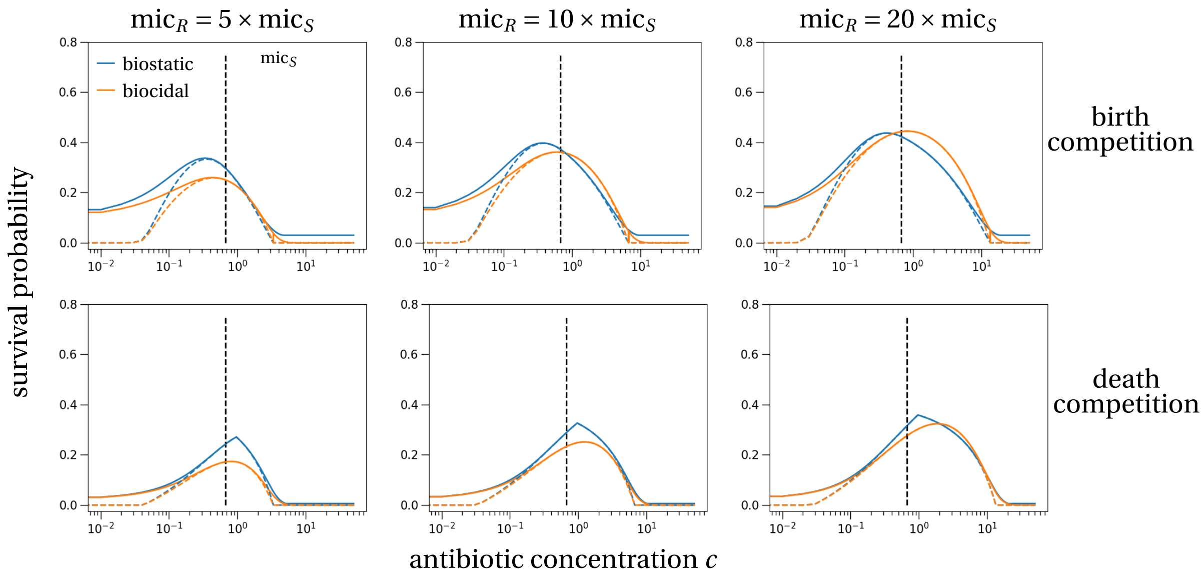


Figure I.3: **Survival probability for the parameterization with tetracycline.**

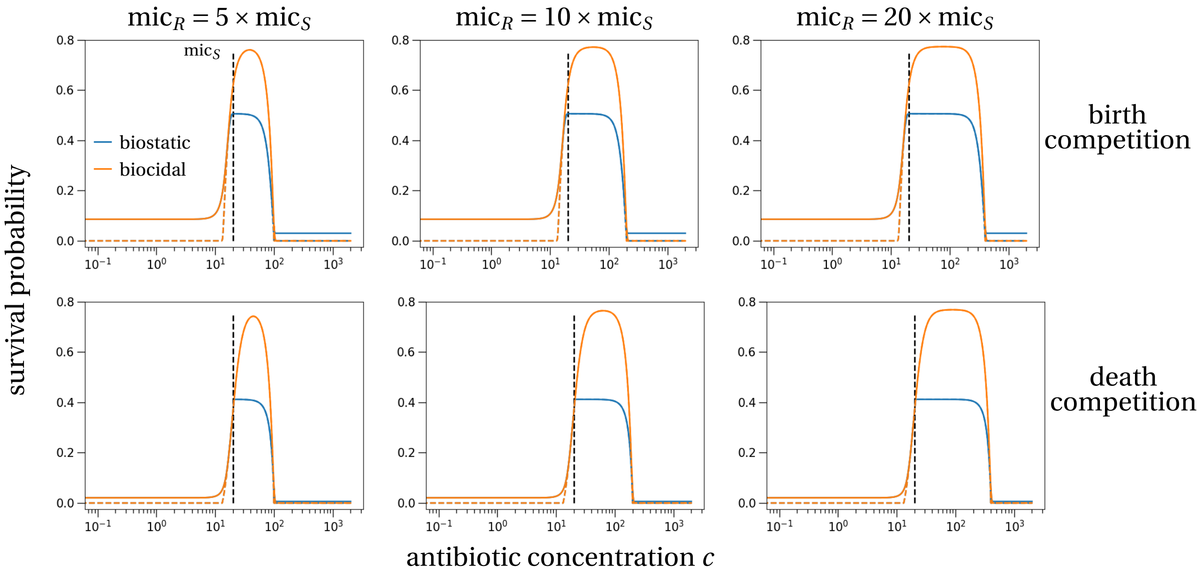


Figure I.4: **Survival probability for the parameterization with hypothetical steep.**

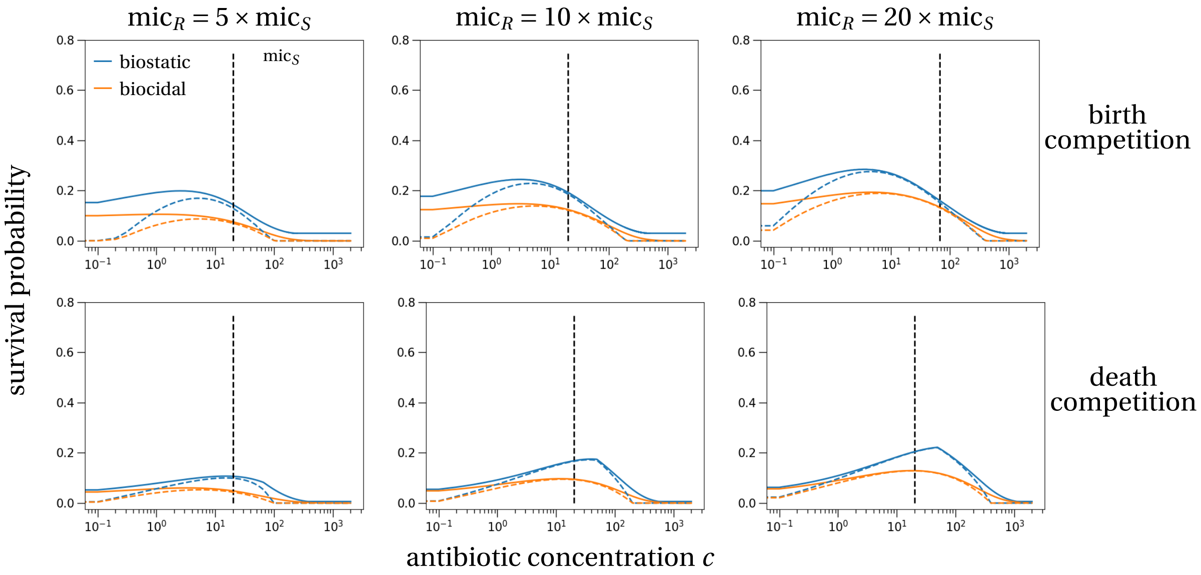


Figure I.5: **Survival probability for the parameterization with hypothetical flat.**

#### I.2 Size at the end of treatment

We also computed the expected size of the resistant subpopulation at the end of treatment for these four alternative parameterizations. Again we find no substantial differences to the main results from the main text: the antibiotic concentration that maximizes the size at the end of treatment is close to the MIC of the sensitive strain (Figs. I.6-I.9).

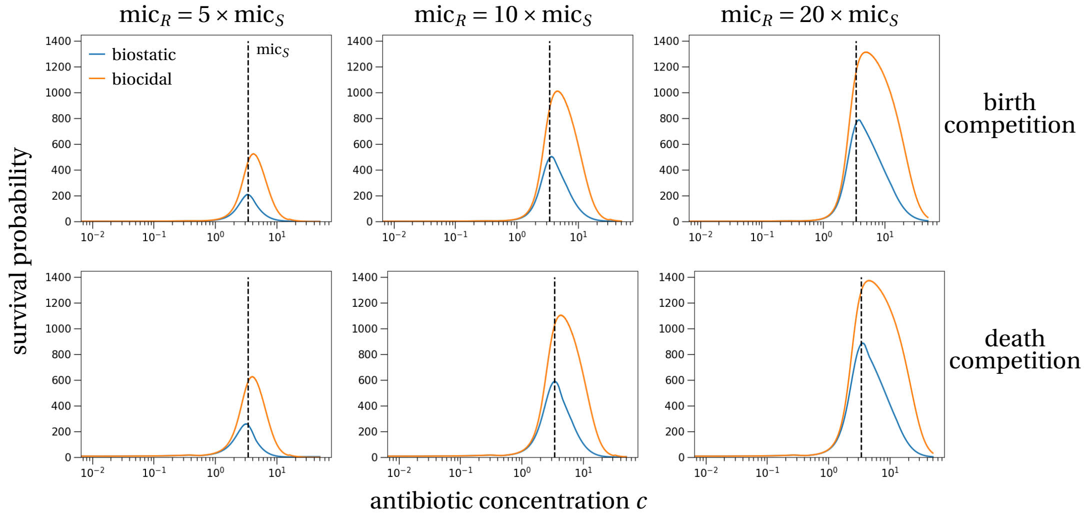


Figure I.6: **Size at the end of treatment for the parameterization with ampicillin.** The solid line shows the theoretical prediction for the size of the resistant subpopulation after a seven day treatment.

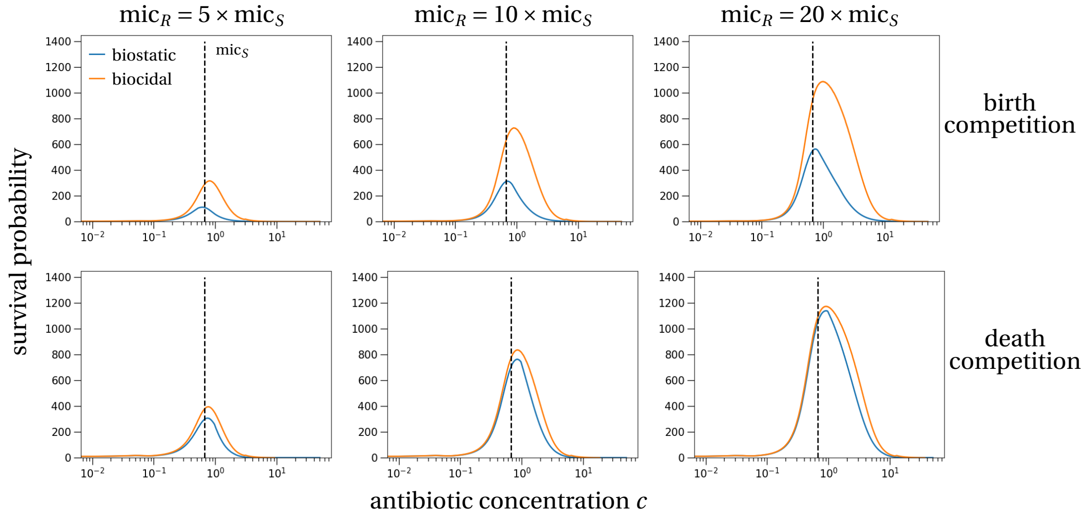


Figure I.7: **Size at the end of treatment for the parameterization with tetracycline.**

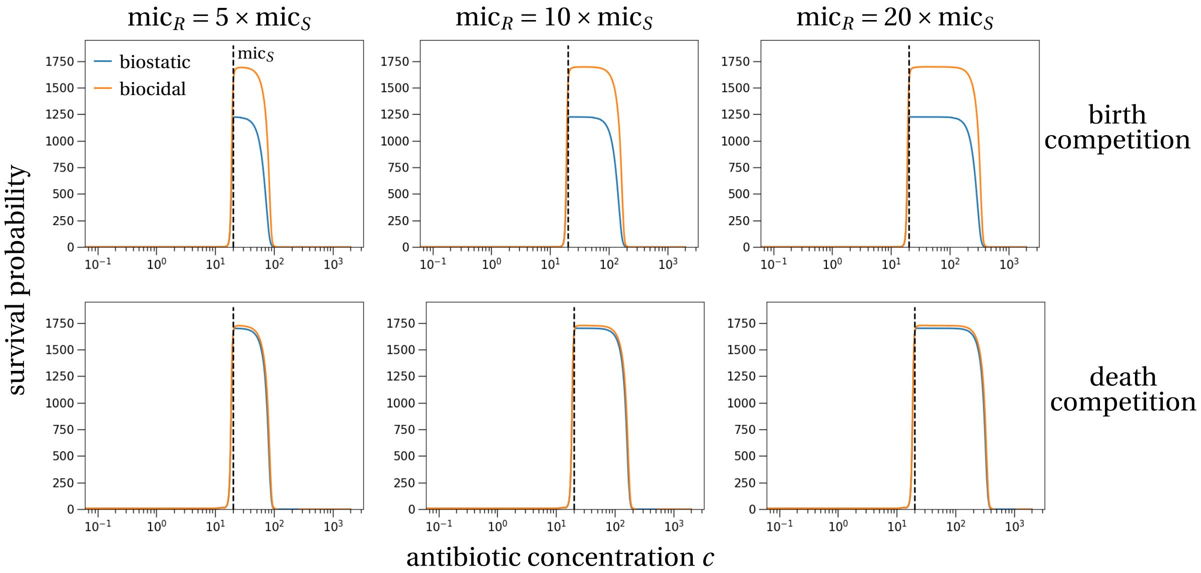


Figure I.8: **Size at the end of treatment for the parameterization with hypothetical steep.**

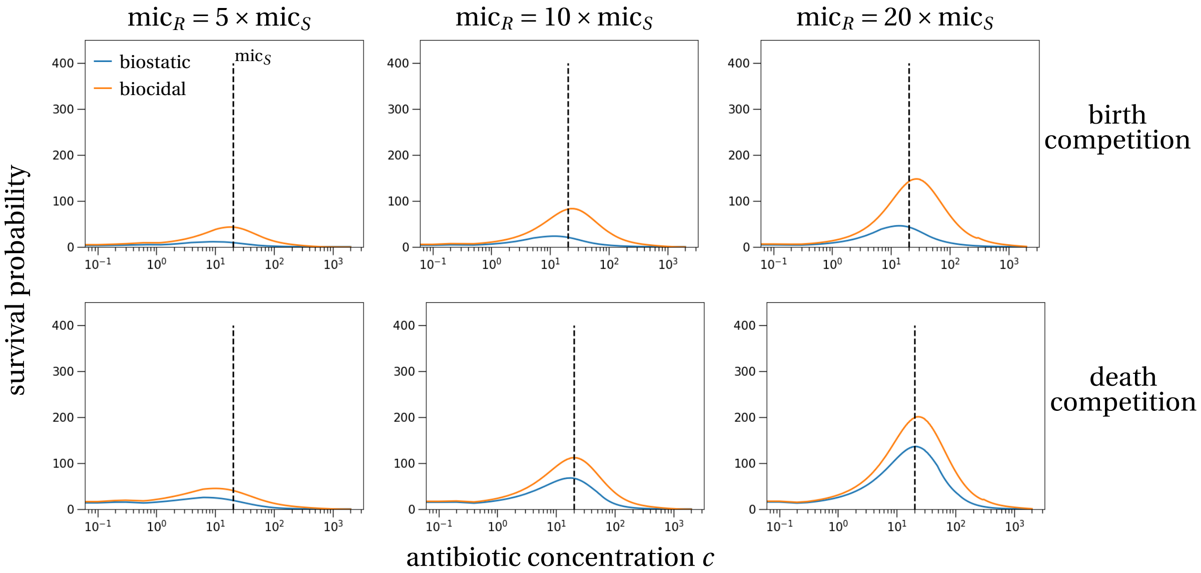


Figure I.9: **Size at the end of treatment for the parameterization with hypothetical flat.**

### J Alternative model with an explicit immune response

In this section, we study an alternative within-host model of antibiotic resistance that explictly includes an immune response. Such a model has previously been studied for example in Day and Read (2016) and Scire et al. (2019).

#### J.1 Model description

Verbally, the model can be described as follows: the sensitive and resistant strains replicate at rates $r_S(c)$ and $r_R(c)$, which under biostatic treatment depend on the antibiotic concentration. Under biocidal treatment these rates are constant. Pathogens are cleared at rates $d_S(c)$ and $d_R(c)$, which depend on the antibiotic concentration in the case of biocidal treatments, otherwise the rates are independent of the concentration. Pathogens trigger an immune response. The production of immune cells at rate $\pi$ scales linearly with the abundance of pathogens. Immune cells, denoted by $I(t)$, degrade at a rate $\mu$. The interaction between immune cells and the pathogen results in the clearance of pathogens at rate $\xi$. For example, in case of biostatic treatment these dynamics result in the following deterministic system:

$$\begin{aligned}\frac{dS}{dt} &= (r_S(c)(1-\eta) - d_S)S(t) - \xi I(t)S(t)\,,\\ \frac{dR}{dt} &= (r_R(c) - d_R)R(t) - \xi I(t)R(t) + \eta r_S(c)S(t)\,,\\ \frac{dI}{dt} &= \pi(S(t)+R(t)) - \mu I(t)\,,\end{aligned} \tag{J.1}$$

where $\eta$ is the mutation rate from the sensitive to the resistant type.

Translated to our model, we have the following pathogen replication and death rates:

$$\begin{aligned} &\text{biostatic:} \qquad r_j(c) = \beta_j - \alpha_j(c), \qquad d_j = \delta_j\,;\\ &\text{biocidal:} \qquad r_j = \beta_j, \qquad d_j(c) = \delta_j + \alpha_j(c)\,. \end{aligned} \tag{J.2}$$

As a comparison, in Day and Read (2016) the following pathogen replication and death rates were used:

$$\begin{aligned} r_S(c) &= 0.6\times(1-\tanh(15(c-0.3))), \qquad d_S = 0.01\,,\\ r_R(c) &= 0.59\times(1-\tanh(15(c-h_R))), \qquad d_R = 0.01\,, \end{aligned} \tag{J.3}$$

where $h_R = 0.45$ in Fig. 2 in and $h_R = 0.6$ in Fig. 3 in Day and Read (2016). This functional form for the effect of an antibiotic on the replication rate results in the same shape of replication rate than our choice of parameterization (Fig. 2 in the main text).

In both models, treatment starts once the patient shows symptoms. We assume, following Day and Read (2016), that the pathogen threshold to develop symptoms is at 100 pathogenic cells. Treatment therefore starts once $S(t)+R(t) \geq 100$. For simplicity, we assume that resistant cells are not present initially, and add a single resistant cell at the onset of treatment (standing genetic variation) or let resistance arise by mutation throughout the simulation (*de novo* emergence).

#### J.2 Sample trajectories

To get an understanding about the within-host dynamics of these different models and assumptions, we plot ten stochastic trajectories for each of the models: our model with resistance by standing genetic variation (Fig. J.1), our model with resistance by de-novo emergence (Fig. J.2), and the model by Day and Read (2016) with de-novo resistance emergence (Fig. J.3).

In our model formulation, we observe that the lowest treatment concentration has the tendency to reliably select for resistance emergence to considerable levels in both cases (standing genetic variation and de-novo). Intermediate treatment concentrations, i.e. between the MICs of the sensitive and resistant strain result in a post-treatment growth of the resistant subpopulation that would be easily detectable so that treatment by another antibiotic could be administered. High antibiotic concentrations above the MIC of the resistant strain suppress the pathogen entirely.

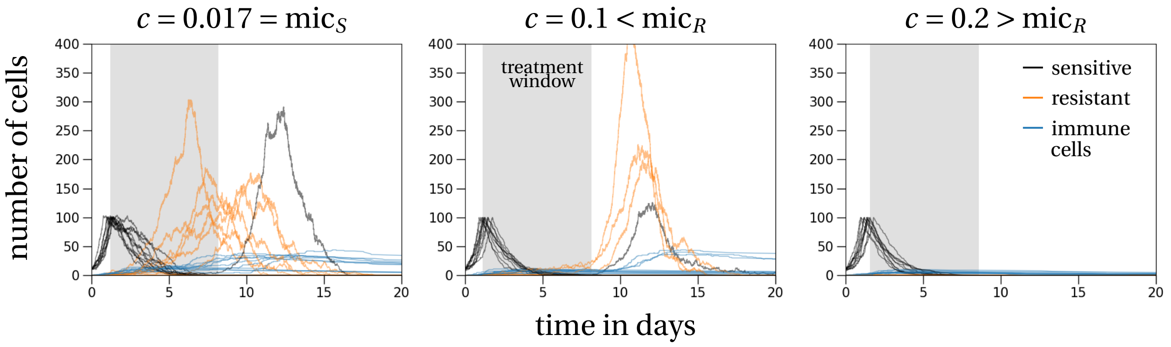


Figure J.1: **Stochastic trajectories of our model with immune response (standing genetic variation).** Ten stochastic trajectories are plotted for different levels of antibiotic treatment with a biostatic drug. Low treatment (left) is at a concentration equal to the MIC of the sensitive strain. Intermediate treatment (middle) corresponds to a concentration just below the MIC of the resistant strain ($\text{mic}_R$=10×$\text{mic}_S$). High concentration (right) has a value just above the MIC of the resistant strain. We assume that resistance arises through standing genetic variation, i.e., we add a single resistant cell at the onset of treatment, which happens when $S(t) = 100$. Treatment then lasts for seven days, depicted for one trajectory by the shaded region. Initial conditions are: $S(0) = 10$, $R(0) = 0$, $I(0) = 0$.

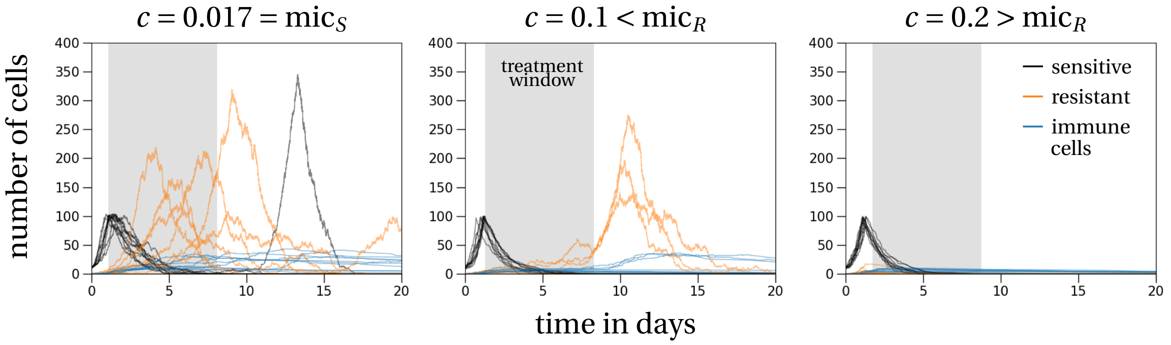


Figure J.2: **Stochastic trajectories of our model with immune response (*de novo* emergence).** The figure has the same structure as Fig. J.1. However, in these simulations resistance arises *de novo* by mutation of sensitive cells at replication. The mutation rate is $\eta = 0.01$. Initial conditions are: $S(0) = 10$, $R(0) = 0$, $I(0) = 0$.

The trajectories obtained from the model implementation by Day and Read (2016) recover the results stated in that paper. However, as stated in the main text, we do not believe that a drug administered at a concentration below the sensitive strain MIC is a realistic scenario to treat an acute disease (left column in Fig. J.3).

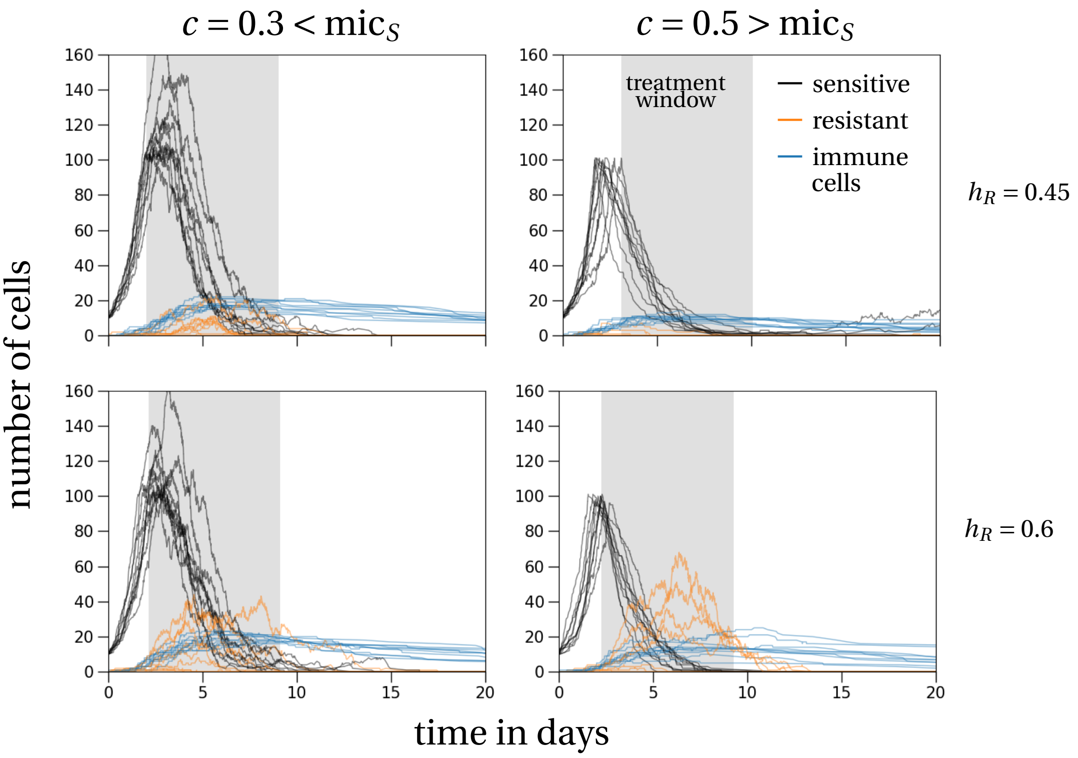


Figure J.3: **Stochastic trajectories of the model by Day and Read (2016) (*de novo* emergence).** Ten stochastic trajectories are plotted for different levels of antibiotic treatment (columns) and different levels of resistance (rows). The top row corresponds to low resistance ($h_R = 0.45$) and the bottom row to high resistance ($h_R = 0.6$). The left column corresponds to sub-MIC treatment ($c = 0.3$) and the right column to an intermediate treatment level ($c = 0.5$). Resistance arises *de novo* by mutation of sensitive cells at replication. The mutation rate is $\eta = 0.01$. Initial conditions are: $S(0) = 10$, $R(0) = 0$, $I(0) = 0$.

#### J.3 Survival probability

Analytical results for the survival probability of the resistant strain seem unattainable in this model because of the complex interaction between immune response and pathogen load, which creates a time-inhomogeneous selection environment that depends on the resistant pathogen load. Therefore, the resistant strain, if it survives, is not just growing exponentially but is limited in growth by its own abundance through the immune response, which impedes analytical progress. We therefore limit the analysis of the survival probability to stochastic simulations.

In the stochastic simulations we have used the default parameter set (Table A.2). To describe the immune response dynamics, we have chosen parameters as in Day and Read (2016): $\xi = 0.075$, $\pi = 0.05$, $\mu = 0.05$. Following our approach of the main text where we study resistance emergence from standing genetic variation, we set the mutation rate equal to zero, $\eta = 0$. The initial number of sensitive and resistant pathogens is $S(0) = 10$, $R(0) = 0$ (we add a resistant cell at the onset of treatment), and the initial number of immune cells is $I(0) = 0$. The survival probability until the end of a seven day treatment is shown in Fig. J.4. As visible, our general finding of a maximal survival probability at the MIC of the sensitive strain translates, at least in this parameter set, to a model with an explicit treatment of immune response.

For a comparison, we also implemented the parameterization of Day and Read (2016). The results

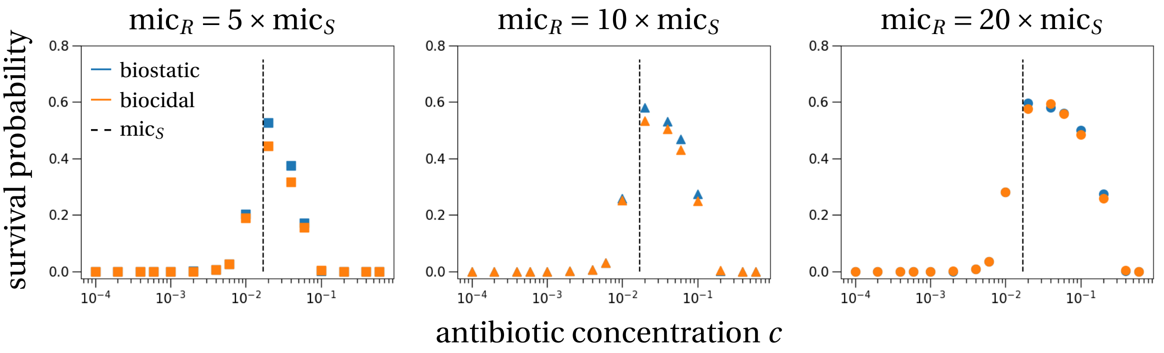


Figure J.4: **Survival probability of the resistant subpopulation in a model with explicit immune response dynamics.** Similar to the model with intraspecific competition, the maximal risk of resistance survival is located at the MIC of the sensitive strain (dashed lines). The MIC of the resistant strain is varied, going from left to right: $5 \times mic_S$, $10 \times mic_S$, $20 \times mic_S$. The difference between the two different drug types, biostatic (blue) and biocidal (orange), is not substantial, at least under this parameterization.

for that model are shown in Fig. J.5 and show that the location of the maximal survival probability of the resistant strain is below the MIC of the sensitive strain. Note that Day and Read (2016) studied *de novo* emergence of resistance, which is what we study here as well. Their parameter choices are: $\eta = 0.01$, $S(0) = 10$, $R(0) = 0$ and $I(0) = 2$, and for the immune response $\xi = 0.075$, $\pi = 0.05$, $\mu = 0.05$.

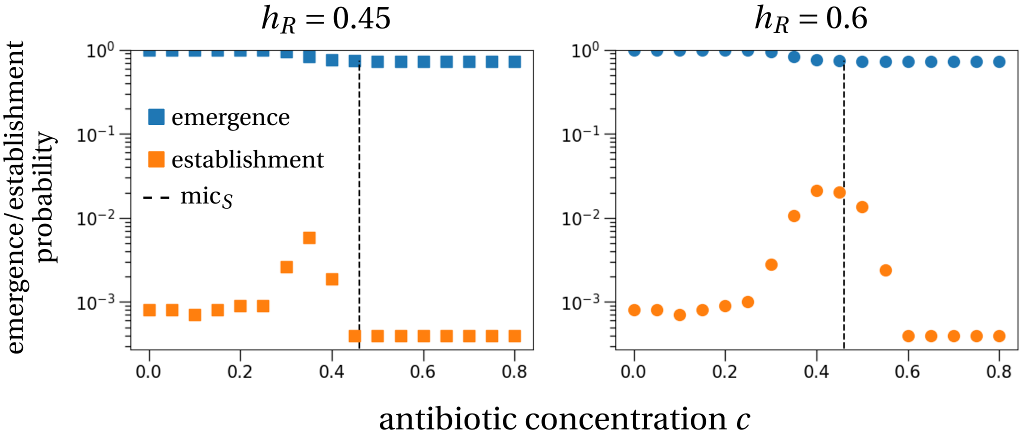


Figure J.5: **Emergence and establishment probability of the resistant subpopulation in the model from Day and Read (2016).** We use the parameterization of Day and Read (2016) as outlined in the text. Emergence (blue symbols) is measured by the appearance of a mutant throughout the 7 days of treatment plus an additional incubation time, i.e. treatment is started once the pathogen population size reaches 100 cells. Establishment (orange symbols) is registered if the resistant subpopulation reaches a size of at least 100 bacterial cells at some point during the incubation period or treatment. The intermediate maximum of the establishment probability is in both cases, low (left) and high (right) level of resistance, again below the MIC of the sensitive strain (dashed vertical lines), which is in line with our previous findings. Note the logarithmic y-axes. Symbols are computed from $10^5$ simulations.

### K Numerical simulations

All stochastic simulations use the exact Gillespie algorithm (Gillespie, 1977). To this end the transition rates need to be specified, which depends on the model at question. We detail the potential events that can happen for biocidal treatment with birth competition and the immune response model from Section J. We assume no mutations in the outline of the models below. If mutations were included they would happen with a mutation probability $\eta$ upon replication of sensitive cells, i.e. with probability $\eta$ a resistant cell is born and with probability $(1-\eta)$ the daughter cell is antibiotic-sensitive.

**Biocidal treatment and birth competition:** Both bacterial types, sensitive and resistant, can have two events: birth or death. These events happen with the following rates ($j$ denotes the bacterial strain):

- *birth* occurs at rate $X_j(t)\left(\max\left(\beta_j - \gamma\frac{X_S(t)+X_R(t)}{K}\right)\right)$, which increases the number of cells of type $j$, $X_j(t)$, by one;
- *death* occurs at rate $X_j(t)(\delta_j + \alpha_j(c))$, which decreases the number of cells of type $j$, $X_j(t)$, by one.

**Immune response model (biocidal drug):** Both bacterial types, sensitive and resistant, can have two events: birth or death. Additionally, the immune response is modeled by the dynamics of immune cells that are produced and degraded. This results in the following four event types (note that in total there are six possible events, because we consider two bacterial strains):

- *bacterial birth* occurs at rate $X_j(t)\beta_j$, which increases the number of cells of type $j$, $X_j(t)$, by one;
- *bacterial death* occurs at rate $X_j(t)(\delta_j + \alpha_j(c) + \xi I(t))$, which decreases the number of cells of type $j$, $X_j(t)$, by one;
- *immune cell birth* occurs at rate $\pi(X_S(t) + X_R(t))$, which increases the number of immune cells, $I(t)$, by one;
- *immune cell death* occurs at rate $\mu\, I(t)$, which decreases the number of immune cells, $I(t)$, by one.

Given any model, the Gillespie algorithm then works as follows (Gillespie, 1977):

1. The population is updated according to one of the possible events. The event is randomly chosen, where the probability for each event to happen is equal to $k/k_{\text{tot}}$, where $k$ is the rate of a specific event and $k_{\text{tot}}$ is the sum of all event rates.
2. The time is updated. A random number from an exponential distribution with mean $1/k_{\text{tot}}$ is drawn and this random number is added to the current time, i.e. $t_{\text{new}} = t_{\text{old}} + dt$, where $dt \sim Exp(1/k_{\text{tot}})$.

The simulations are run until an end condition is met, which depends on the purpose of the simulation, e.g. end of treatment for the survival probability or size of the resistant population size at the end of treatment, clearance of resistance for the carriage time, etc.
